# Multiplex single-cell chemical genomics reveals the kinase dependence of the response to targeted therapy

**DOI:** 10.1101/2023.03.10.531983

**Authors:** José L. McFaline-Figueroa, Sanjay Srivatsan, Andrew J. Hill, Molly Gasperini, Dana L. Jackson, Lauren Saunders, Silvia Domcke, Samuel G. Regalado, Paul Lazarchuck, Sarai Alvarez, Raymond J. Monnat, Jay Shendure, Cole Trapnell

## Abstract

Chemical genetic screens are a powerful tool for exploring how cancer cells’ response to drugs is shaped by their mutations, yet they lack a molecular view of the contribution of individual genes to the response to exposure. Here, we present sci-Plex-**G**ene-by-**E**nvironment (sci-Plex-**G**x**E**), a platform for combined single-cell genetic and chemical screening at scale. We highlight the advantages of large-scale, unbiased screening by defining the contribution of each of 522 human kinases to the response of glioblastoma to different drugs designed to abrogate signaling from the receptor tyrosine kinase pathway. In total, we probed 14,121 gene-by-environment combinations across 1,052,205 single-cell transcriptomes. We identify an expression signature characteristic of compensatory adaptive signaling regulated in a MEK/MAPK-dependent manner. Further analyses aimed at preventing adaptation revealed promising combination therapies, including dual MEK and CDC7/CDK9 or NF-kB inhibitors, as potent means of preventing transcriptional adaptation of glioblastoma to targeted therapy.

## Introduction

The response of individual cancer cells to treatment with a drug depends on myriad factors, including but not limited to their locations within a tumor, proximity to vessels, epigenetic histories, and, of course, their genotypes. Defining the contribution of individual genetic mutations to how a tumor will respond to a given drug regimen is critically important to personalized therapy and for understanding cancer pathobiology. However, dissecting the mechanisms by which each of the many genes frequently mutated in cancer confers drug resistance is extremely challenging because the space of gene-drug interactions is enormous. Addressing this requires a scalable, systematic approach for quantifying a drug’s effect on a cell of a given genotype. Chemical genetics, the study of how exogenous compound exposures interact with cells and alter gene-product function and phenotype ^1^, is a powerful means to define the genetic dependencies on treatment response across genetically distinct samples. The induction of genetic heterogeneity via targeted genetic perturbation ^2–4^ (e.g., CRISPR/Cas9) has drastically increased the power of such screens allowing for the systematic determination of how perturbed gene activity alters response. However, these screens are largely limited to determining the effect of the genetic perturbation on gross phenotypic outcomes (viability, cell growth) or very specific molecular readouts (reporter expression, enzymatic activity). Moreover, CRISPR-based chemical genetic screens with simple phenotypic readouts are largely applied at the population level. Therefore, they are unable to link genotype to cellular response in a precise, mechanistic manner. There is therefore a need for novel methods by which to systematically interrogate the genetic requirements of effective drug treatment.

Single-cell CRISPR screens ^5–8^ allow in-depth molecular insight into the effects of genetic perturbation of genes associated with diverse biological processes, including those prioritized by bulk CRISPR screens ^9,10^. Recently, single-cell CRISPR screens have been performed at genome-scale, providing a rich map of the effect of perturbation of all expressed genes ^11^. To further probe genetic dependencies to exposure using a single-cell genomic readout necessitates incorporating additional strategies to multiplex at the level of exposure (drugs, doses) within an experiment. This multiplexing allows the assay to scale to large combinatorial spaces and minimizes batch effects due to sample processing. Recently, we developed sci-Plex, a nuclear hashing approach that couples high-throughput chemical screens to combinatorial indexing RNA-seq (sci-RNA-seq3) ^12^, allowing for the analysis of the molecular effect of thousands of chemical perturbations in parallel ^13^. Here, we present sci-Plex gene-by-environment or sci-Plex-GxE, which extends sci-Plex to pooled single-cell CRISPR screens, markedly increasing the number of unique gene-exposure interactions tested within one experiment and providing the opportunity to define how large cohorts of genes affect the response of cells to many exposures.

As an initial proof-of-principle, we apply our approach to probe the relationship between exposure to the standard-of-care alkylating agent temozolomide (TMZ) ^14^ and genetic perturbation of the mismatch repair (MMR) pathway, a known genetic dependency to S_N_1 alkylating agent-induced damage ^15,16^. Using this system, we develop computational methods for determining the extent to which a genetic perturbation interferes with or enhances the expected effects of a drug on the transcriptome. We then apply sci-Plex-GxE to determine the molecular consequence of genetic perturbation of the 522 kinases in the human protein kinome ^17^ on the response of 3 glioblastoma (GBM) cell lines to 4 small molecules inhibitors targeting the receptor tyrosine kinase pathway, the most frequently over-activated pathway in GBM ^18^ and a driver for glioma initiation ^19^ and maintenance ^20^. We find that drug exposure leads to the induction of a transcriptional program characterized by the upregulation of genes capable of eliciting an adaptive (i.e., drug-induced) resistance phenotype ^21,22^. Our single-cell genetic screen prioritized kinases involved with the regulation of MAPK, replication, cell division, and stress signaling that modulate the expression of this adaptive compensatory program. Combinatorial chemical exposure targeting of a subset of these kinases confirmed their involvement in regulating this transcriptional response and co-treatments that limit the ability of a cell to mount an adaptive response to kinase therapy.

## Results

### sci-Plex-GxE combines nuclear hashing and CRISPR-based single-cell genetic screens

To determine the contribution of individual genes to the response to chemical exposure at scale, we combined our single-cell chemical transcriptomics platform ^13^ with the CROP-seq system for single-cell CRISPR/Cas9 genetic screens ^8^. We developed and optimized a method for the enrichment of sgRNA containing transcripts ^5,6,23^ from within the context of sci-RNA-seq3. Our enrichment strategy relies on targeted capture of the CROP-seq derived sgRNA containing puromycin transcript in combination with standard poly-A mRNA capture during RT and amplifying sgRNA containing transcripts from the final sci-RNA-seq3 mRNA library (**Fig. 1A & Methods**).

**Figure 1:**
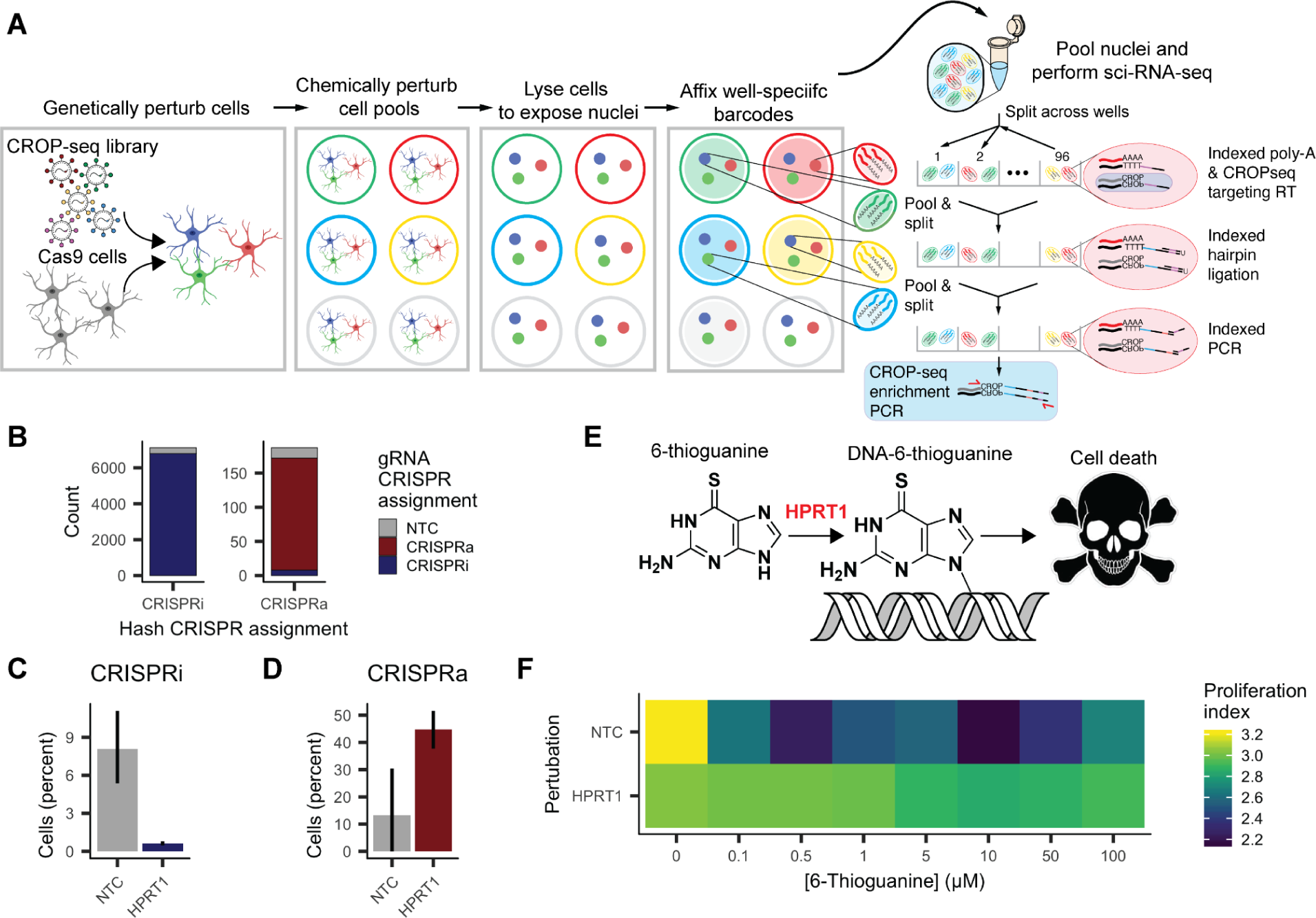
sci-Plex-GxE couples genetic and perturbation screens with high sensitivity and specificity. **A**) Overview of sci-Plex-GxE workflow including targeted enrichment of CROP-seq derived gRNA containing transcripts. **B**) Concordance between CRISPRi and CRISPRa gRNA assignments derived from the hash added to individually labeled CRIPSRi and CRIPSRa pools (hash CRISPR assignment) and by the sequence of captured gRNAs (gRNA CRISPR assignment). **C/D**) Percent of cells expressing *HPRT1* in cells expressing systems for CRISPR-mediated knockdown (**C**) or overexpression (**D**) and gRNAs targeting *HPRT1* or NTC controls. **F**) Aggregate expression of genes associated with proliferation in NTC and *HPRT1* knockdown cells after exposure to 6-thioguanine.

To determine the specificity and sensitivity of our assay, we performed a sgRNA cell mixing experiment. We transduced A172 GBM cells expressing either dCas9-KRAB for gene knockdown (CRISPRi) or dCas9-SunTag for gene overexpression (CRISPRa) ^24^ with CROP-seq-OPTI libraries containing either optimized CRISPRi or CRISPRa sgRNAs ^25^ targeting the *HPRT1* locus, a modulator of cell sensitivity to the chemotherapeutic agent 6-thioguanine (6-TG), and non-targeting controls (NTCs). We arrayed CRISPRi and CRISPRa perturbed cell pools across columns of a 96-well plate and exposed cells to increasing concentrations of the purine analog 6-TG or DMSO control. After 96 hours, cells in individual wells were harvested and subjected to our sci-Plex GxE protocol (**Fig. 1A** and **Methods**). We captured 18,585 single-cell transcriptomes and used our sci-Plex hash labels to remove doublets and to confidently assign one well/treatment condition to 17,549 cells (94.4%). We identified one or more sgRNAs in 15,589 of these treatment-assigned singlet cells (88.8%) (**Supp Fig. 1A**), of which 94.4% expressed 1 sgRNA at a high proportion (**Supp Fig. 1B-C**), consistent with the low multiplicity of infection of our transduction. We next compared cell assignment according to captured sgRNAs or sci-Plex hashes. As expected, cells containing hashes denoting CRISPRi wells were largely assigned a CRISPRi sgRNA and vice-versa (**Fig. 1B**). Cells expressing CRISPRi and CRISPRa sgRNAs against *HPRT1* displayed a decrease and increase in *HPRT1* expression, respectively (**Fig. 1C, 1D**). Loss of HPRT1 activity leads to resistance to the nucleic acid analog 6-thioguanine (6-TG) ^26^ by decreasing its incorporation into DNA (**Fig. 1E**). As expected, *HPRT1* knockdown cells had a high expression of genes associated with proliferation in the presence of increasing doses of 6-TG compared to NTC controls (**Fig. 1F**). This experiment confirms that sci-Plex-GxE can detect a genetic requirement for individual cells’ response to a drug exposure via global transcriptome analysis.

### A chemical genomics approach to prioritize genotypes with strong effects on the response of cells to exposure

The scalability and multiplexing ability of sci-Plex-GxE at the level of genotypes and exposures allows profiling of the effects of gene-exposure interactions at scale. However, there are additional considerations for its application in chemical genomic screening, namely a need for analysis workflows that allow prioritization of genotypes within large-scale perturbation screens and a way to summarize complex transcriptional effects compactly. To arrive at analytical solutions to these challenges, we first applied our approach to a known genetic dependency to alkylation damage.

Temozolomide (TMZ) is an oral alkylating agent and the standard-of-care for glioblastoma brain cancer chemotherapy ^14^, whose toxicity is mediated by the formation of *O^6^*-meG lesions in the DNA. A cytotoxic response to TMZ primarily depends on the expression of methyl guanine methyltransferase (MGMT) and functional DNA mismatch repair (MMR) (**Fig. 2A**), with the activity of these pathways mediating resistance and sensitivity, respectively ^27^. We profiled the transcriptional consequence of exposing MMR-perturbed A172 CRISPRi cells to increasing doses of TMZ for 96 hours. We targeted MMR pathway components that comprise the *O^6^*-meG recognition complex (MutSɑ: *MSH2* and *MSH6*), the MMR processing complex downstream of lesion recognition (MutLɑ: *MLH1* and *PMS2*) (**Fig. 2A**), and controls including targeting of an MMR component not involved in the processing of *O^6^*-meG (*MSH3*), *MGMT,* which is epigenetically silenced in A172 ^28^ and non-targeting controls (**Supp Fig. 2C-D**).

**Figure 2:**
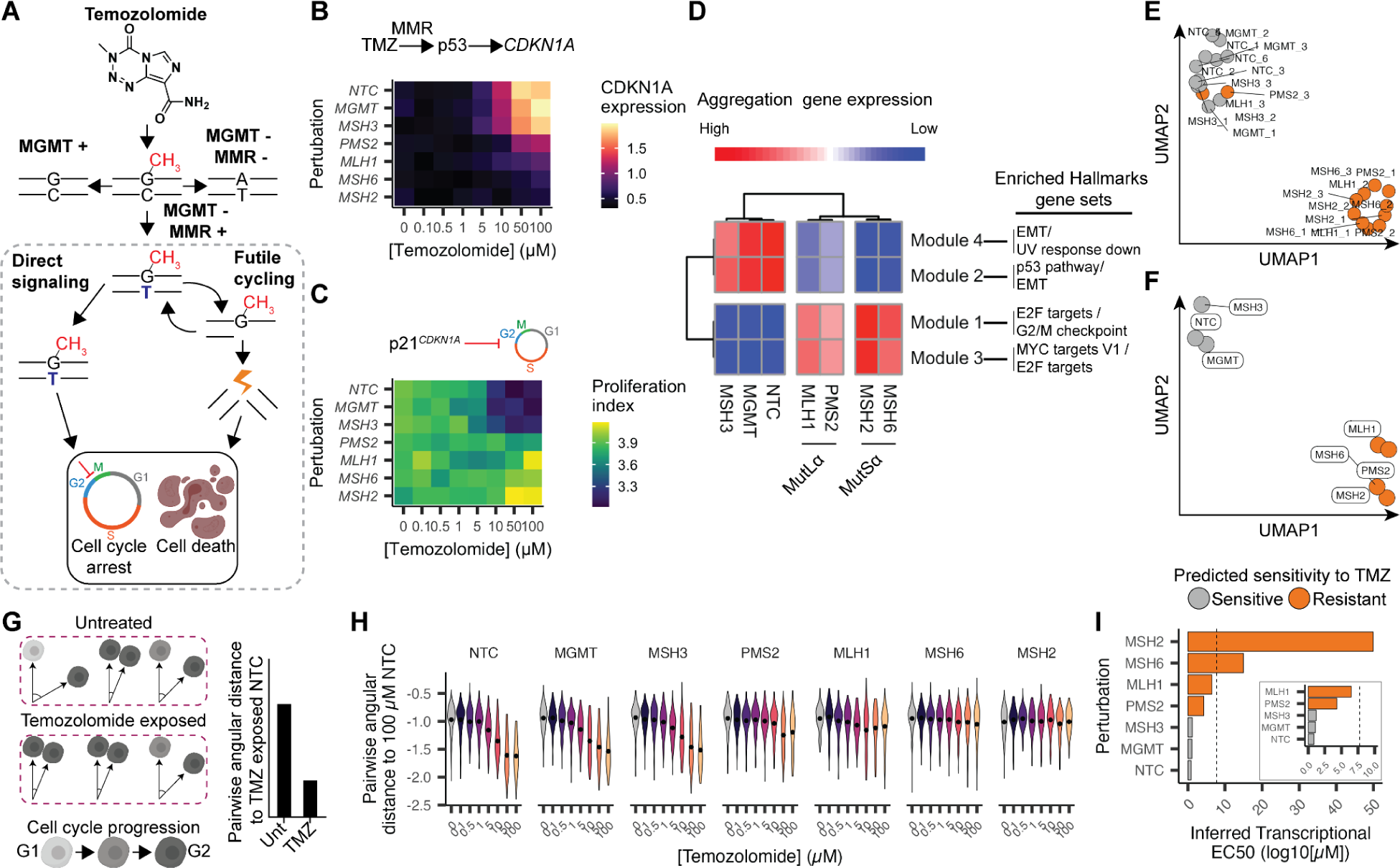
Defining the relationship between genotypes and summarizing the magnitude of the perturbation after combined genetic and chemical perturbation. **A**) Mechanism of TMZ-induced mismatch repair dependent *O*^6^-methyguanine toxicity. **B**) Heatmap depicting *CDKN1A* expression levels as a function of perturbation via CRISPRi-mediated knockdown and exposure to TMZ. **C**) Heatmap depicting the aggregate expression of genes associated with proliferation as a function of perturbation via CRISPRi-mediated knockdown and exposure to TMZ. **D**) Left: Heatmap depicting the aggregate expression of gene modules derived from genes that are differentially expressed as a function of genotype in TMZ-exposed and genetically perturbed cells. Expression was aggregated for cells exposed to high doses of TMZ (10, 50, and 100 µM). Right: Enriched MSigDB Hallmarks gene sets for gene modules in the experiment. **E/F**) UMAP embedding derived from the proportions of cells expressing individual gRNAs (**E**) or gRNAs against the labeled target (**F**) across PCA clusters in our experiment. **G**) Cartoon depicting the expected decrease in angular distance as cells enact a transcriptional response and associated cell cycle arrest after TMZ exposure. **H**) Violin plots depicting the pairwise angular distance of every cell to the mean expression of NTC cells exposed to 100 µM TMZ for all genotypes after exposure to increasing doses of TMZ. **I**) Inferred transcriptional effective concentration (TC50) is defined as the concentration of drug necessary to reach 50% of the change in angular distance exhibited by TMZ-exposed NTC cells. Dashed line: maximum molarity of an aqueous solution as a threshold for genotypes where the drug cannot induce 50% of the effect observed in NTC. Insert excludes MSH2 and MSH6.

Cells expressing sgRNAs against *MGMT*, *MSH3,* or NTC controls displayed a robust, dose-dependent increase in the expression of the cell cycle inhibitor and p53 target *CDKN1A* (**Fig. 2B**) that was accompanied by decreases in the expression of genes associated with proliferation (**Fig. 2C**). Analysis of differentially expressed genes (DEGs, FDR < 10%) revealed a strong correlation (Pearson’s rho 0.50-0.94) across genotypes expected to alter sensitivity to TMZ at the highest doses of drug (**Supp Fig. 2F**). The magnitude of these changes was decreased in cells expressing sgRNAs against *MLH1* and *PMS2* and largely abrogated in cells expressing sgRNAs against *MSH2* and *MSH6* (**Fig. 2B-C**). Defining gene modules across the union of DEGs recovered signatures that define genotypes sensitive to TMZ exposure (NTC, *MGMT,* and *MSH3*) and further subdivided genotypes associated with mismatch recognition (*MSH2* and *MSH6*) and downstream processing (*MLH1* and *PMS2*) (**Fig. 2D**). Of note, gene modules associated with p53 signaling (module 2) varied depending on the perturbed MMR complex consistent with activation of DNA damage signaling by lesion recognition before MutLɑ processing ^29^.

We next sought to summarize differential responses to exposure. Dimensionality reduction did not identify unique cellular states induced by the interaction of genotypes and TMZ (**Supp Fig. 3A-D**), likely due to the fact that phenotypes perturbed by modulation of MMR activity (e.g., proliferation, cell cycle arrest) are available to non-perturbed cells. However, MMR perturbations did alter the distribution of cells across these shared phenotypes (**Supp Fig. 3E**), which could be summarized by dimensionality reduction techniques (**Fig. 2E-F**). Although this approach can prioritize genotypes within the context of a screen, its reliance on defining cell states a priori is limiting. Therefore, we sought an approach tailored to prioritizing genotypes in the context of the response to drug exposure.

**Figure 3:**
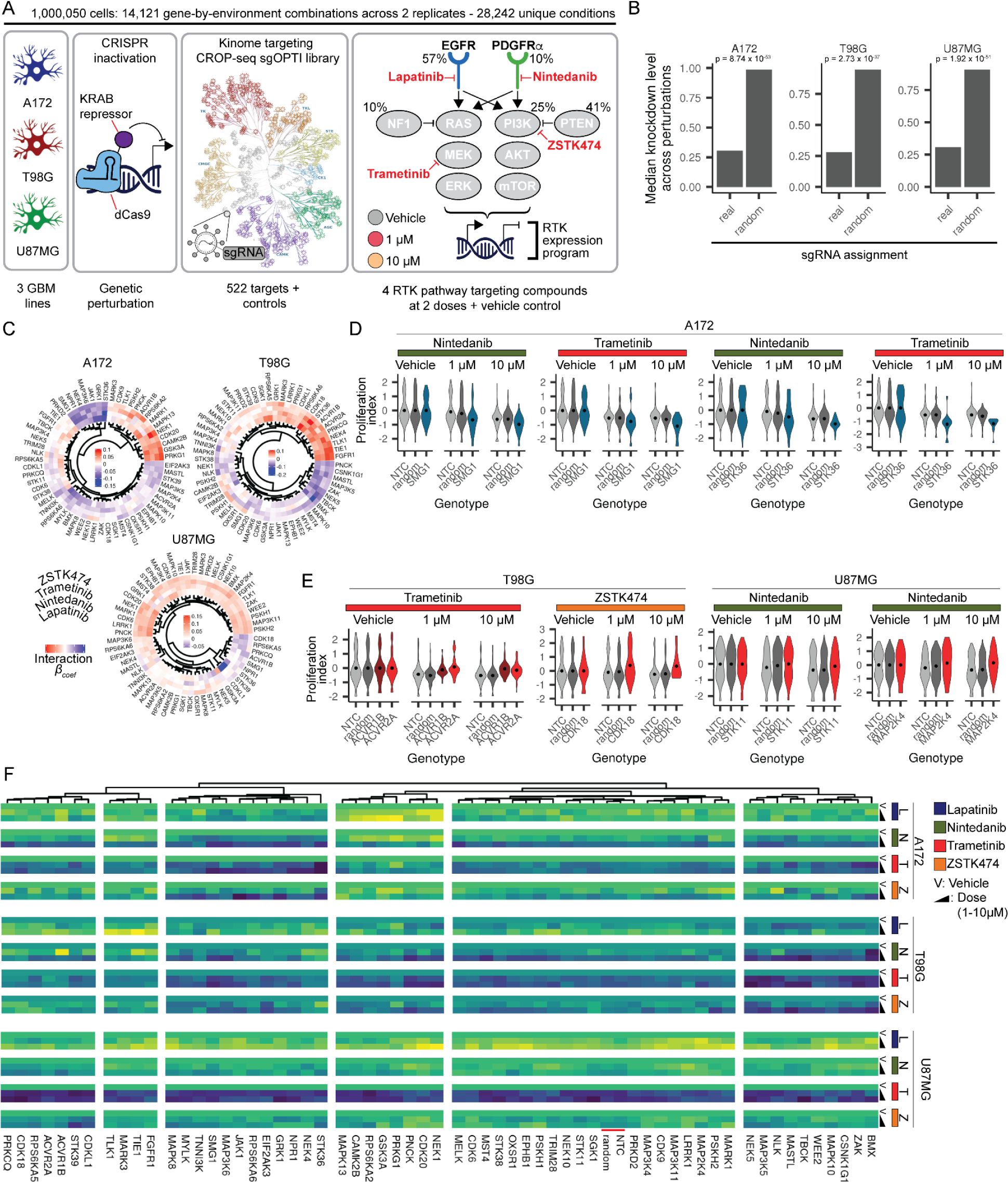
A single-cell kinome targeting genetic screen identifies subtle effects of perturbation on proliferation-associated gene expression. **A**) Schematic depicting the sci-Plex-GxE screen to determine the contribution of the kinome to the transcriptional response of glioblastoma cells to RTK pathway targeted therapies. **B**) Median knockdown level across the three cell lines in our screen as a function of sgRNA assignment (real) or a random permutation of sgRNA assignment labels (random) (Wilcoxon rank sum test). **C**) Hierarchical clustering of β coefficients for the term from a generalized linear model describing the interaction between drug treatment and kinase perturbation on the aggregate expression of proliferation-associated genes across kinase perturbations with a significant interaction term in response to at least one treatment in one cell line. **D**) Heatmap depicting the mean aggregate expression of proliferation-associated genes for proliferation perturbing and controls genotypes. Values were centered as in B/C. Only genotypes with more than 5 cells at top doses are shown. The red annotation bar highlights non-targeting and random targeting control genotypes. **E/F**) Violin plots depicting the aggregate expression of proliferation-associated genes for the cell lines, treatment, and genotypes presented. For each genotype, values were centered on the mean of untreated cells.

We devised a strategy to describe how close a given cell is to the expected response to a given drug, identifying gene-by-environment interactions in a given condition. We first identified a set of genes that are dynamically regulated as a function of drug exposure in “wild-type” NTC cells (**Supp Fig. 3F**). Second, for every dose, we calculated the pairwise transcriptome distance for every cell relative to the averaged expression profile of NTC cells exposed to the highest dose of drug based on this set of drug-responsive genes (**Supp Fig. 3G**). We then quantify the extent to which a perturbation deviates from the change in pairwise transcriptome distance of unperturbed cells as they converge on a drug-induced phenotype, in this case, a TMZ-induced cell cycle arrest (**Fig. 2G**). TMZ exposure led to a decrease in pairwise transcriptome distance across unperturbed NTC cells and our negative control knockdowns (*MGMT* and *MSH3*). Whereas knockdown of *PMS2*, *MLH1*, *MSH6,* and *MSH2* altered the dose-response relationship in pairwise transcriptome distance to NTC as a function of dose (**Fig. 2H & Supp. Fig. 3H-J**).

We used this framework to define a transcriptional effective concentration 50 or “TC50” ^13^ to determine, for each genotype, the concentration of drug necessary to arrive at 50% of the transcriptional response observed in NTC cells. TC50s were similar for NTC, *MGMT* and *MSH3* knockdowns, whereas loss of *MLH1*, *PMS2*, *MSH6* or *MSH2* renders cells largely less sensitive to drug (**Fig. 2I & Supp. Fig. 3K-L**) as previously described ^30^. We find our transcriptome distance approach to be robust to the number of cells per genotype (**Supp. Fig. 3M**), able to detect a graded dose-dependent response to a drug, and able to quantify the progression of a given cell along that response, characteristics necessary for identifying gene-by-environment interactions within the context of large-scale single-cell chemical genomic screens.

### Defining the contribution of the human protein kinome to the transcriptional response induced by kinase inhibition

Having demonstrated the ability of sci-Plex-GxE to detect the genetic requirements for exposure to drugs, we sought to systematically characterize the genes that determine a tumor’s response to standard-of-care therapy at scale. Glioblastoma brain cancer is characterized by overactive receptor tyrosine kinase (RTK) signaling, with ∼90% of tumors presenting with an activating mutation in the RTK pathway ^18,31^. Paradoxically, GBM patients display low response rates to RTK-targeted therapy despite these prominent alterations in RTK signaling. Adaptive ^32^ (i.e., pharmacologically-induced) activation of pathways that rescue RTK signaling and/or induce similar downstream effectors is suspected to be amongst the most common mechanisms by which tumors evade therapy ^21,22,33,34^, including GBM ^35,36^. Mapping out how tumors shift their gene expression programs to evade the detrimental effects of drug exposure could identify new opportunities to improve therapy.

To determine the contribution of an entire class of genes to drug-induced transcriptional adaptation ^37,38^ in GBM, we perturbed all members of the human protein kinome ^17^ in 3 GBM cell lines expressing the dCas9-KRAB CRISPRi system. Heterogeneous cell pools were then exposed to one of four compounds targeting the receptor tyrosine kinases EGFR (lapatinib) and PDGFRɑ (nintedanib), and the MAPK and PI3K signaling pathway components MEK (trametinib) and PI3K (zstk474) at two doses (1 and 10 µM) or vehicle control for 72 hours and subjected to sci-Plex-GxE. Our screen contained 14,121 gene-by-environment combinations across two independent transductions for 28,242 unique conditions across 1,052,205 single-cell transcriptomes (**Fig. 3A, Supp. Fig. 4A**). After excluding putative doublets, we assigned a condition to 991,940 single-cell transcriptomes (94.3% of cells) and observed good agreement in expression across replicates in our experiment (**Supp. Fig. 4B**). We identified a sgRNA in 988,276 cells and a median target knockdown of ∼70% across our panel of targeted kinases (**Fig. 3B**). We note that we observed low sgRNA assignment from our sgRNA PCR enrichment strategy applied to a small amount of our final mRNA libraries (data not shown). The addition of multiple rounds of sgRNA enrichment PCR increased our assignment rate suggesting the low assignment from 1 or a few enrichment cycles is due to a bottleneck when amplifying from large, complex libraries.

**Figure 4:**
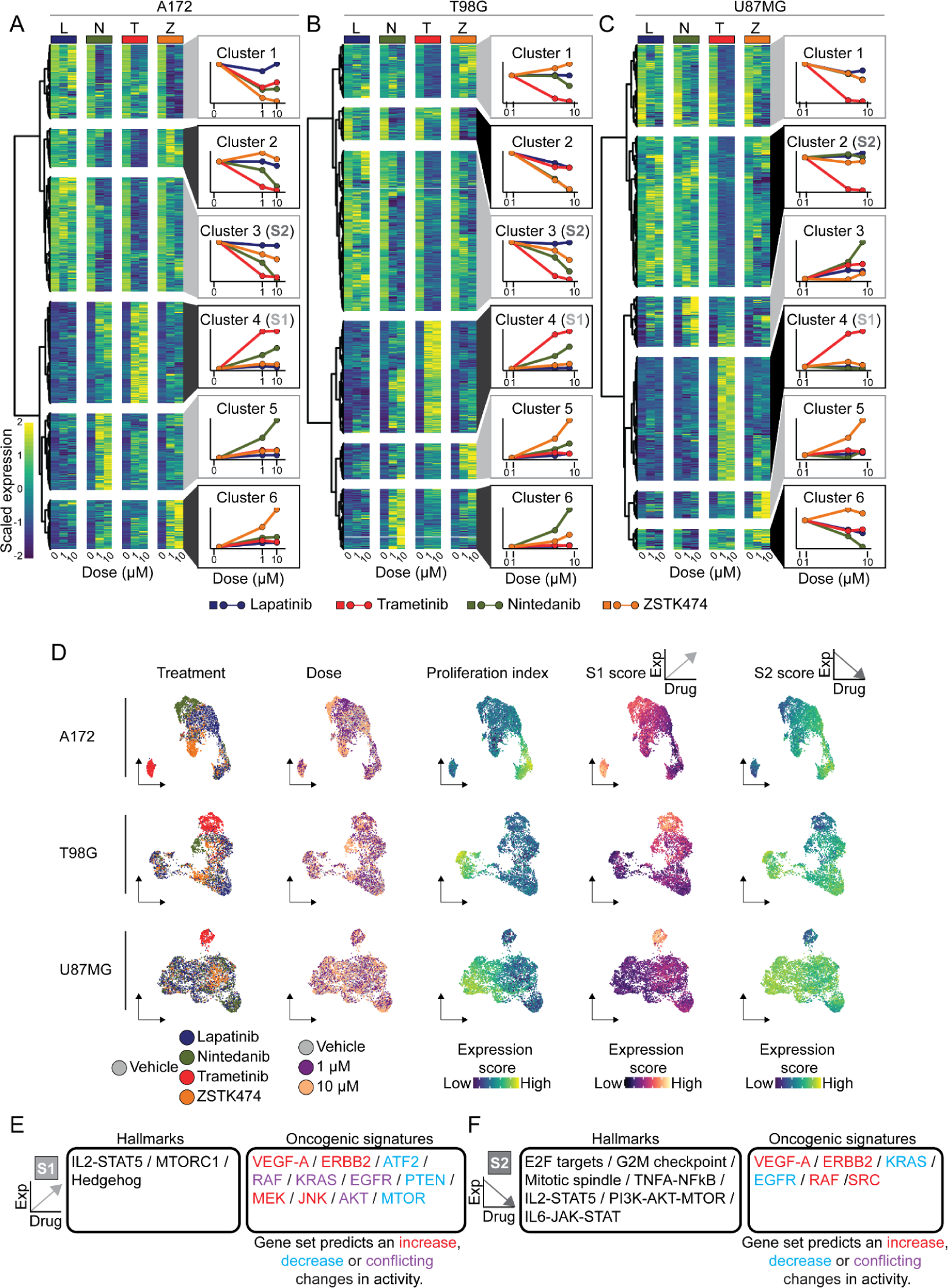
Exposure to small molecules targeting the RTK pathway drives changes in gene expression, including dynamic alterations in kinase expression. **A-C**) Heatmaps depicting the average expression of genes that are dynamic as a function of exposure to at least 1 of 4 small molecules targeting components of the RTK pathway in A172 (**A**), T98G (**B**) and U87MG (**C**) cells (FDR < 5% & |β_coef_| > 1). Right panels: Aggregate gene expression across clusters for gene clusters to the left for A172 (**A**), T98G (**B**), and U87MG (**C**) cells. Colors denote the individual treatments as in the top annotations in panels **A-C**. **D**) UMAP embeddings of unperturbed A172 (top), T98G (middle), and U87MG (bottom) cells colored by treatment, dose, proliferation index, or aggregate scores for the conserved upregulated S1 or downregulated S2 signatures. Arrows denote the dynamics of S1 and S2 signature expression as a function of RTK pathway inhibition. **E-F**) Gene set enrichment analysis using the MSigDB hallmark, and oncogenic signatures gene set collections of signatures that increase (**E.** S1) or decrease (**F.** S2) in expression as a function of drug exposure with RTK targeted therapy. Arrows as in **D**.

Examining the proportion of genotypes in vehicle-exposed cells to the starting plasmid proportion revealed a depletion of gRNAs targeting kinases that are likely required by all three cancer cell lines. We observed the strongest depletion for 16 kinases across one or more cell lines (|z-score| > 2, **Supp Fig 5A**). These included kinases involved in mitosis (*AURKA*, *AURKB*, *BUB1B*, *PLK1*) ^39^ and ribosome maturation (*RIOK1* and *RIOK2*) ^40,41^. We did not identify kinases that conferred a similarly strong growth advantage across the cell lines in our study when knocked down.

**Figure 5:**
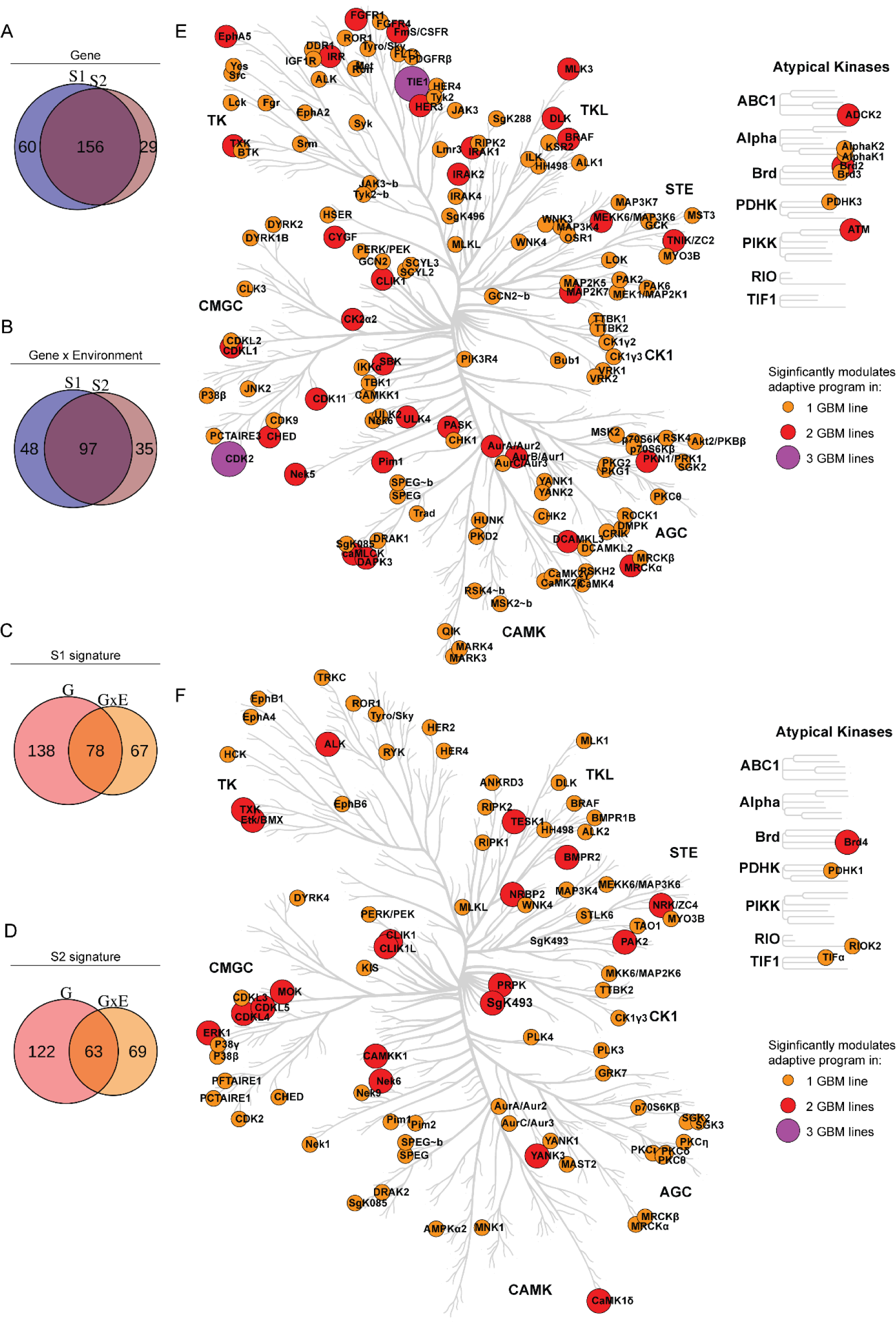
Perturbation of individual kinases alters the global transcriptional response to RTK pathway targeting small molecules. **A-B**. Venn diagram of the overlap between kinases whose knockdown leads to a significant shift in the S1 or S2 signatures of the putative adaptive program without (**A**) or with (**B**) a significant interaction effect. **C-D**. Venn diagram of the overlap between kinases whose knockdown leads to a significant shift in the expression of the S1 (**C**) or S2 (**D**) signatures of the putative adaptive program without or with a significant interaction effect. **E**. Human protein kinome tree, adapted from CORAL^64^, highlighting kinases whose knockdown leads to a significant shift in the expression of a putative adaptive program. The size and color of each circle denote whether we identified a significant effect in 1, 2, or all GBM cell lines (FDR < 5%). **F**. Kinome tree as in **A** highlighting kinases whose knockdown leads to a significant shift in the expression of a putative adaptive program for which we identify a significant interaction effect with drug exposure. The size and color of each circle denote whether we identified a significant effect in 1, 2, or all GBM cell lines (FDR < 5%).

### sci-Plex-GxE reveals kinases required for the transcriptional response to inhibiting the RTK pathway

We next sought to define the transcriptional changes induced in NTC cells by targeting the RTK pathway with small molecules. Exposure to compounds targeting the RTK pathway decreased cell viability, with the strongest effect observed for cells exposed to trametinib and the weakest effect for lapatinib (**Supp Fig 5B**). This decrease in cell viability was accompanied by a decrease in the expression of genes associated with proliferation (NTC unperturbed genotype, **Fig 3C-F**).

To identify kinases required for maintaining expression of the proliferation gene program, we next modeled their expression as a function of drug, dose, genotype, and their interaction using generalized linear models ^12^. These models included kinase-by-drug “interaction terms”, which capture effects observed in drug-treated, genetically perturbed cells that are not observed in vehicle-treated, genetically perturbed cells or NTC cells exposed to a drug. We identified 60 kinases involved in a significant interaction term for at least one exposure. We observed concordance for interaction terms across treatments within each cell line (**Fig 3**) and high similarity in proliferative expression profiles across our controls (**Fig 3D**).

Our analysis identified kinases that led to a significant decrease in proliferative gene expression across multiple treatments. For example, in A172 cells, knockdown of the genotoxic stress response PI3K-like kinase *SMG1* ^42^ or the positive regulator of hedgehog signaling *STK36* ^43^ led to a significant decrease in proliferation in response to both nintedanib and trametinib (**Fig 3E**), suggesting these kinases are required for proliferation in cells treated with these drugs. We also identified kinases whose knockdown led to increased proliferative gene expression. These included the *ACVR1B ^44^*, *ACVR2A ^45^*, *STK11* (LKB1) ^46^, and *MAP2K4* ^47^ tumor suppressors (**Fig 3F**). Knockdown of *CDK18*, recently described as a co-factor for ATR-driven homologous recombination repair in GBM ^48^, led to a significant increase in proliferation in response to the PI3K inhibitor zstk474 (**Fig 3F**). Zstk474, like other PI3K inhibitors, targets other DNA damage response kinases such as DNA-PKc and ATM ^49,50^ and was shown to generate strand breaks in GBM cells ^50^. Therefore, the effect of CDK18 loss on the proliferative response to zstk474 exposure may result from an additive increase in genotoxic stress. Our analysis demonstrates that our multiplex chemical genomic screen identifies significant interactions between genotype and exposure, including kinase perturbations that sensitize or resist the effect of RTK pathway targeting inhibitors on proliferative gene expression. However, because sci-Plex-GxE profiles the entire transcriptome, it is not limited to viability or proliferation phenotypes and in principle could characterize the genetic requirements of other gene programs, including transcriptional adaptation to targeted therapy.

### Single-cell RNA-seq identifies shared transcriptional responses to inhibition of receptor tyrosine kinase signaling in glioblastoma cell lines

Targeting over-activated oncogenic kinases induces drastic remodeling of gene expression networks ^21,22,33,34^, enabling tumors to substitute an alternative pathway to restore signaling, a process termed adaptive resistance ^51^. To quantify transcriptional adaptation in our GBM lines, we performed regression analysis of the effects of each drug on each gene’s expression using generalized linear models. We identified robust dose-dependent changes in expression with 4,553, 3,112, and 3,149 genes differentially expressed after exposure to 1 or more inhibitors in A172, T98G, and U87MG cells, respectively (**Fig. 4A-C & Supp. Fig. 6A-C**). We observed strong transcriptional effects upon trametinib exposure and modest changes in cells exposed to lapatinib, consistent with their effects on cell viability (**Supp Fig 4A**). Comparing drug-induced transcriptional responses revealed a large overlap in the genes altered in the three lines when exposed to the FGFR/VEGFR/PDGFR family inhibitor nintedanib and trametinib or zstk474 (**Supp Fig 6D-F**), suggesting that RTKs of the FGFR/VEGFR/PDGFR families, are largely responsible for driving MEK and PI3K activity in these cell lines.

**Figure 6:**
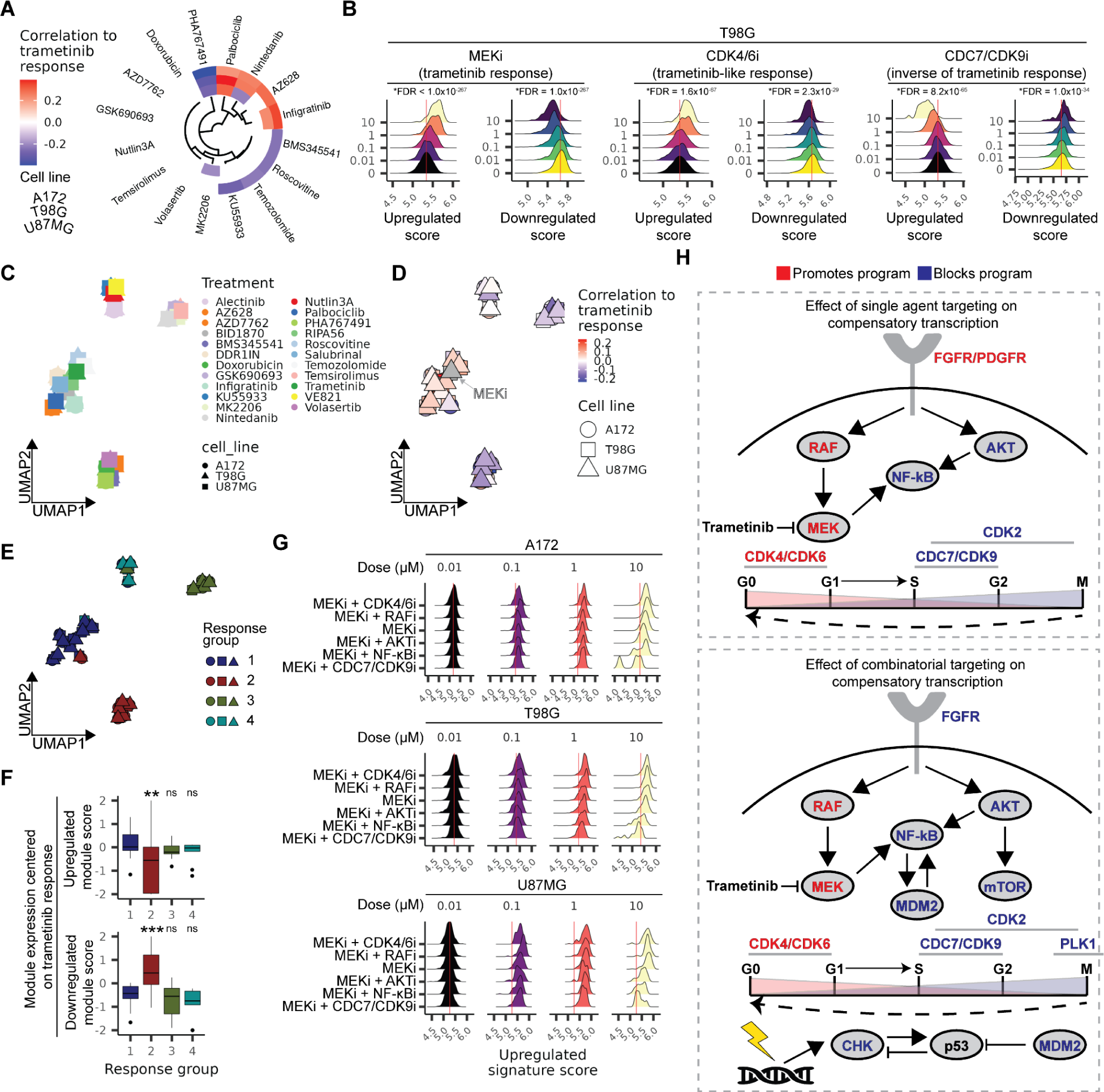
Single and combinatorial kinase inhibition identifies chemical regulators of MEK inhibition-dependent dynamic expression changes. **A**) Circos heatmap of the correlation of specified single drug exposures to the compensatory program enacted by MEK inhibition with trametinib. Only significant correlations (FDR < 0.05) over a cutoff of Pearson’s ⍴ ±0.2 are shown. **B**) Density plots of upregulated and downregulated signature scores of T98G cells treated with the MEK inhibitor trametinib (MEKi), the CDK4/6 inhibitor palbociclib (CDK4/6i) or the CDC7 inhibitor PHA767491 (CDC7i) for the three glioblastoma cell lines. Red vertical lines denote the mean signature expression of vehicle-exposed cells. *FDR < 0.05. **C**) UMAP embeddings summarizing the pair-wise correlation of all specified combinatorial trametinib exposures across genes that compose the compensatory program enacted by RTK pathway inhibition. Shapes refer to individual cell lines. **D**) UMAP as in **C,** colored by the correlation of each combinatorial trametinib exposure to cells exposed to trametinib alone across genes that compose the compensatory program enacted by RTK pathway inhibition. **E**) UMAP as in **C** colored by response group clusters identified by Leiden-based community detection. **F**) Boxplots of the expression of signatures that make up the compensatory program. Values for each combinatorial exposure are centered on the response of cells to trametinib alone, such that combinations that increase signature expression relative to trametinib are positive and vice-versa for those that decrease signature expression relative to trametinib. **G**) Density plots of upregulated signature scores of GBM cells treated with the MEK inhibitor trametinib (MEKi) alone or in combination with the RAF inhibitor AZ628, the AKT inhibitor GSK690693, teh NF-kB inhibitor BMS345541, and the dual CDC7/CDK9 inhibitor PHA767491. Plots are ordered from top to bottom by the effect of treatment on the signature. **H**) Pathway summary of proteins whose targeting alone (**top**) or in combination with MEK inhibition (**bottom**) blocks (blue) or exacerbates (red) the compensatory program enacted by MEK kinase inhibition.

We next sought kinases that were themselves altered at the RNA level in response to drug exposure, as these might potentiate regulation of the broader adaptive response. Hierarchical clustering of all differentially expressed genes identified modules of genes with similar dose-dependent changes across our treatments (**Fig. 4A-C**), including 156, 121, and 126 kinases whose expression was significantly altered in response to exposure in a cell line- and drug-specific manner. Exposure of A172 and U87MG cells to nintedanib led to a pronounced increase in *ERBB4* expression, whereas trametinib exposure led to a significant increase in *ERBB4* in A172 (**Supp Fig. 6G**). T98G did not display a significant increase in *ERBB4* expression upon exposure to trametinib, but we did observe a strong increase in *EPHA5* (**Supp Fig. 6G**). Other kinases displayed similar responses to a given treatment across all cell lines. For example, exposure of GBM cell lines to trametinib resulted in a significant decrease in the expression of *WEE1*, a kinase that negatively regulates the mitotic kinase CDK1/CDC2 ^52^ (**Supp Fig. 6G**).

To identify genes with a response to each drug shared across cell lines, we calculated the Jaccard index to investigate the number of genes shared between each drug-induced gene module across every pairwise set of cell lines (**Supp Fig. 6H-K)**. We identified two sets of genes with similar dynamics as a function of drug exposure across the 3 cell lines, one of which broadly increases with dose (“S1”), and another that decreases (“S2”) (**Fig. 4A-C**, **Fig. 4D** and **Supp Fig. 6L**). The genes in these shared drug-induced modules also had largely concordant responses to RTK pathway inhibition in a panel of 4 patient-derived GBM lines, indicating they may constitute a core program of GBM transcriptional response to RTK pathway targeting ^53–55^ (**Supp. Fig. 6M-N**).

To assess the extent to which genes in the core adaptive program are known targets of cancer-associated signaling pathways, we performed gene set enrichment analysis (GSA) (**Fig 4E-F**). The downregulated S2 gene module was enriched for genes associated with the regulation of the cell cycle (**Fig 4F**), consistent with a decrease in pro-proliferative signaling downstream of RTK pathway inhibition. However, S1 and S2 genes did not neatly map onto gene sets known to be upregulated or downregulated in response to inhibiting the RTK pathway, respectively. For example, the down-regulated S2 module was enriched for genes that report on active KRAS and PI3K-AKT-MTOR signaling, with their decrease in expression suggesting a block of these pathways. In contrast, the up-regulated S1 module was enriched for genes associated with active mTORC1 signaling, which may report on the activation of a distinct subset of the mTORC1 program (**Fig. 4D-E**). Similarly, mixed results were obtained using the MSigDB oncogenic signatures gene set collection ^56,57^, where modules displayed enrichment for gene sets that suggest activation and inactivation of different subsets of the RTK signaling pathway (**Fig 4D-E**, right panels). To identify genes that may mediate an escape to RTK pathway inhibition, we further examined the list of genes that make up the S1 upregulated drug-induced module. This revealed inhibitor-induced increases in the expression of kinases central to activation of the RTK pathway, including the receptor tyrosine kinase *EGFR*, the cytoplasmic tyrosine kinase *ABL1,* the dual specificity kinase *MAP2K1,* which encodes the ERK activator MEK1, and the PI3K catalytic subunits *PIK3C2A* and *PIK3C2B*. Together, these observations are consistent with the induction of a complex drug and cell-line-specific adaptive, compensatory program with a shared core component that may mediate survival in response to RTK pathway inhibition.

In order to identify kinases that are required for transcriptional adaptation in each cell line, we quantified how perturbation of each kinase alters the expression of the core adaptive gene modules. We used our pairwise transcriptome distance framework to identify kinases whose loss leads to a deviation in the drug-induced expression of S1 and S2, finding a significant effect due to perturbation of 55, 97, and 84 kinases in A172, T98G, and U87MG cells, respectively. We identified a high overlap between kinases that regulate the S1 and S2 gene modules in the absence of an interaction with drug (gene effects) or with a significant interaction between genotype and drug (gene x environment effects). In all, we identified 156 kinases whose perturbation altered compensatory program expression at the level of gene effect (**Fig. 5A**, FDR < 5%) and 97 kinases that altered compensatory program expression with evidence of a gene x environment effect (**Fig. 5B**, FDR < 5%). In contrast, comparing the list of kinase modulators within a given gene module revealed low overlap between kinases with significant gene and gene x environment effects (**Fig. 5C-D**).

Across our set of kinase hits, we identified 42 kinases with a significant gene effect (**Fig. 5E**) and 23 with a significant gene x environment effect (**Fig. 5F**) in 2 or more GBM cell lines. Amongst hits, only *CDK2* and *TIE1* significantly affected the compensatory program across all 3 GBM lines. CDK2 activity is critical for progression along the late-G1 and early-S phase of the cell cycle, is frequently overactivated in various cancers due to the upregulation of its binding partner cyclin E ^58^, and contributes to adaptation to CDK4/6 inhibition ^59^. *TIE1* encodes an orphan receptor tyrosine kinase most frequently associated with endothelial cells and the regulation of angiogenesis by modulating the activity of the TIE2 RTK ^60^. In cancer, TIE2 protein expression has been identified outside of the endothelial compartment and is positively correlated with increased tumor grade in glioma ^61^. Moreover, *TIE1* expression has been shown to induce resistance to chemotherapy in ovarian cancer by modulation the expression of DNA damage repair proteins through activation of the transcription factor KLF5 ^62^.

Amongst kinases with a significant interaction effect is *BRD4* (**Fig. 5A**), an epigenetic reader of histone acetylation and atypical kinase that phosphorylates the c-terminal domain of RNA polymerase II and serves as a master regulator of eukaryotic transcription ^63^. Previous studies have shown that BRD4 activity mediates adaptive transcriptional resistance to MEK inhibition in triple-negative breast cancer ^21^. Our results suggest that BRD4 serves a similar role in response to RTK pathway targeting in GBM cells.

In all, the kinases required for compensation are involved in diverse cellular processes, including cell cycle progression (*AURKA*, *AURKB*, *BUB1*, *PLK3*, *PLK4*, *VRK1, VRK2*), ribosome maturation (RIOK2), cytoskeletal reorganization (*CDC42BPA*, *CDC42BPB*) and proliferative MAPK signaling (*ALK*, *AKT2*, *BRAF*, *ERBB2, ERBB3, ERBB4, FGFR1, MAP4K1, MAPK3*, *SRC*). Given the enrichment for cell cycle processes, we explored whether transcriptional changes upon targeting the RTK pathway are a consequence of accumulation at a particular cell cycle stage as opposed to a response to cellular stress. We observed a correlation between proliferation, G1/S, and G2/M scores in untreated cells and our signatures (**Supp Fig. 7A-B**). However, exposing cells to RTK pathway inhibitors led to altered expression of these genes regardless of proliferation or inferred cell cycle stage (**Supp Fig. 7C-D**), suggesting that the cell cycle stage, baseline RTK pathway activity, and possibly their interaction, can affect this adaptive compensatory program.

### sci-Plex-GxE identifies kinases required for transcriptional adaptation to targeted therapy

A major goal of our workflow is to define the genes required for escaping a therapy, which might then suggest targets for new combinatorial therapies with better efficacy. Therefore, we sought to recapitulate the effects of CRISPR-based knockdown on individual kinases using small molecules. We exposed cells to one of 23 compounds (Supp Table 3) targeting the activity of 16 kinase hits from our screen (AKT, ALK, ATM, RAF, CDK, CHEK, DDR, EIF2AK, FGFR, IKK [CHUK], MEK, PDGFR, PLK, RIPK, RPS6K families) prioritized from those that are hits in more than one cell line or for which related kinases are hits in one cell line (CHEK1, CHEK2) as well as those that are directly involved in RTK signaling (AKT). We also exposed cells to 3 compounds targeting kinases in closely related pathways (ATR, CDC7, CDK4/6, MTOR) and small molecules that produce or are involved in response to genotoxic cell stress (temozolomide, doxorubicin, p53 activator) alone or in combination with the MEK inhibitor and potent inducer of the compensatory program trametinib (**Fig. 5B**). After 72 hours, cells were harvested and unique conditions multiplexed using sci-Plex and subjected to sci3-RNA-seq capturing 213,404 nuclei across single- and combinatorial-drug exposures.

We first investigated the ability of each chemical in isolation to phenocopy the response to trametinib exposure. We performed a correlation analysis of the expression of gene signatures between cells treated with trametinib, the strongest inducer of these signatures, and compounds that had a measurable effect on trametinib-regulated transcription as monotherapy (defined as compounds with > 100 DEGs in two or more cell lines, FDR < 5%). Exposure to the CDK4/6 inhibitor palbociclib elicited the strongest “trametinib-like” response across all cell lines (**Fig. 6A-B**, p < 0.05 and Pearson’s rho > +/-0.2 and **Supp Fig. 8A**). The next highest trametinib-like responses were elicited by inhibition of the RTKs PDGFR (nintedanib) and FGFR (infigratinib) and the inhibition of its upstream regulator RAF (AZ628), although this varied by cell line (**Fig. 6A, Supp. Fig. 8B-D**). Exposure to the dual CDC7/CDK9 inhibitor PHA767491 was strongly anti-correlated to the effects of trametinib exposure (**Fig. 6A-B**). CDC7, or DBF4-dependent kinase (DDK), promotes replication initiation by phosphorylating the minichromosome maintenance (MCM) helicase complex ^65^. CDK9 complexes with cyclin T to form the positive transcriptional factor elongator (pTEFb) complex, a regulator of RNA polymerase II ^66^. The role of CDK9 suggests the possibility that altered regulation of at least the subset of the program upregulated by exposure is a result of a non-specific effect on global transcription. However, the concomitant increase in the downregulated signature (**Fig 6B**) suggests that the observed effect of PHA767491 on drug-induced transcription is not due to a non-specific effect and may be due to additional roles for CDK9 in progression across the S phase of the cell cycle ^67,68^. Although *CDK9* is a hit in our screen, we cannot rule out that additional loss of CDC7 activity and its effect on the cell cycle do not contribute to this drug-induced phenotype.

We next sought to identify combinatorial exposures of drugs that could block the adaptive program induced by trametinib alone. We used generalized linear regression to model the effects of co-exposure on each gene in the adaptive program. We then calculated the correlation of gene-level effects across all co-exposures, grouped combinatorial treatments by the similarity of transcriptional effects summarized across this correlation space, and visualized the results with UMAP (**Fig. 6C**). We identified 4 groupings of combinatorial exposures, including those that differed by the extent of induction of the trametinib induced signature as defined as the correlation to trametinib exposure alone (**Fig. 6D-E**).

Response group 2 had the largest anti-correlated effect to trametinib exposure alone. Response group 2 was composed of co-exposures with AZD7762 (CHKi), BMS345541 (NF-kBi), doxorubicin (topoisomerase IIi), PHA767491 (CDC7/CDK9i), and volasertib (PLK1i) across all 3 cell lines and infigratinib (FGFRi) for 2 of the 3 cell lines and the group significantly attenuated induction of the compensatory program (**Fig. 6F**, FDR < 1%). Response group 3, made up of co-exposures with GSK690693 (AKTi), MK2206 (AKTi), temsirolimus (MTORi), nintedanib (PDGFRi) in all 3 cell lines, and Nutlin3A (MDM2i) in 2 of 3 cell lines was also anti-correlated to trametinib exposure alone. However, the average effect of the response group on the aggregated expression of compensatory modules was not significantly different from trametinib alone. AKT and mTOR have previously been identified to enact compensatory signaling ^69^, and this group may reflect modest blocks to the compensatory program that are not evident across the full signature. In A172 cells, we also found evidence that co-exposure with the RAF inhibitor AZ628 and the CDK4/6i palbociclib exacerbated the compensatory program (**Fig. 6G**). This may have implications for the emergence of resistance to BRAF-mutated tumors treated with combinatorial MEK and RAF inhibition ^70–72^.

Interestingly, these differential responses to combinatorial inhibition could not be readily explained based on differential effects on viability. For example, trametinib co-exposure with palbociclib and PHA767491 had similar dose-dependent effects on viability despite opposing effects on compensatory program expression (**Supp. Fig. 9**). Our combined chemical and genetic perturbation screen defined a compensatory transcriptional response to MEK inhibition and identified kinases that significantly regulate the two gene modules that compose this program. Our chemical genomics approach validated the dependence of this program on CDK4/6 activity and demonstrated that the inhibition of several kinases, including CHK, CDC7/CDK9, FGFR, IKK, AKT, and mTOR interfered with the ability of GBM cells to mount this compensatory program (**Fig. 6H**).

## Discussion

Defining the molecular basis by which individual genes alter the response to therapy has important implications for cancer treatment. The multiplexing ability afforded by single-cell screens is particularly well-suited to probe the large combinatorial space of genetic perturbations and drug treatments. Here we introduce sci-Plex-GxE, a workflow for high throughput chemical genomic screens at single-cell resolution, and demonstrate its capability to identify the genetic architecture that drives response to exposure by investigating the effect of the human protein kinome on the response to RTK pathway inhibition in glioblastoma tumor cells.

The rapid advance of cancer genomics has identified genetic variants and mutations that provide cells with the capacity for malignant transformation and the acquisition of key phenotypes that define a tumorigenic cell state ^73^. Genetic screens have arisen as a powerful means to identify cancer dependencies ^3,4,74^ as well as regulators of toxicity in response to therapeutic exposure ^75^. However, most of these screens report on a limited set of phenotypes (e.g. proliferation or viability). They cannot discern, for example, whether dependencies that similarly alter viability differentially induce unwanted secondary effects that can result in the development of resistance.

Perturb-seq approaches that combine CRISPR-Cas gene editing with a single-cell transcriptomic readout have been applied at genome-scale, defining the effect of genetic perturbation of all expressed genes on transcriptional networks ^11^. Our recent development of multiplexing approaches that allow single-cell technologies to be used in high-throughput chemical transcriptomics screens and its combination with Perturb-seq methods provide an opportunity to understand how cancer cells respond to therapy at scale. To meet this need, we developed sci-Plex-GxE, a platform for multiplex genetic and chemical perturbation screens at single-cell resolution. We demonstrate that our method remains sensitive to capturing both cell-expressing and exogenous tags that report on genetic perturbation (gRNA-containing transcripts) and chemical exposure (cell hashing) while scaling to genome-wide contexts. Further, we developed a computational workflow to prioritize genotypes that significantly shift drug-induced gene expression changes efficiently.

To demonstrate the ability of our approach to identify the genetic requirements of the response to exposure, we applied sci-Plex-GxE to define the contribution of all kinases in the human protein kinome to the dynamic response of glioblastoma cells to the inhibition of 4 nodes in the RTK pathway, all frequently overactivated in the disease. We identified diverse molecular changes in response to RTK pathway inhibition with our genetic screen revealing many kinases whose loss significantly alters proliferative gene expression, suggesting an increased sensitivity to detect growth changes compared to bulk CRISPR screening. We also identified two transcriptional modules whose expression changes are conserved across all three of the cell lines screened, which we posit are part of one conserved transcriptional program. This program is associated with changes in the expression of components of the RTK pathway, including evidence of an adaptive resistance program characterized by increased expression of genes that can activate or bypass RTK signaling. In all, our genetic screen identified 213 kinases whose loss led to significant differences in the induction of this program. We used a chemical genomic approach to validate the contribution of a subset of these kinases to the regulation of this putative adaptive program. We identified compounds targeting cellular activities that can positively (CDK4/6i, FGFRi, PDGFRi, RAFi) and negatively (AKTi, IKKi, CDK2, CDC7/CDK9i) modulate this adaptive program in isolation (**Fig. 6H**, top panel). In addition, we identified compounds that significantly modulate the induction of this adaptive program in the presence of its activation via trametinib exposure. In particular, we find that combinatorial inhibition of MEK kinase and either of AKT, CDK2, CDC7/CDK9, FGFR, MTOR, MDM2, NF-kB, and PLK signaling can block the induction of the core adaptive transcriptional program in GBM cells (**Fig. 6H**, bottom panel). These combinations may be promising combinatorial therapies that minimize the induction of unwanted resistance-associated changes in response to MEKi monotherapy. However, our study is limited to the prioritization of inhibitor combinations based on a desired transcriptional effect. In-depth biochemical and *in vivo* functional characterization are necessary to confirm the ability of combinatorial exposures to increase progression-free survival.

Interestingly, our validation experiment targeting FGFR activity had opposing effects as mono or combination therapy, which may highlight context dependence of transcriptional adaptation in GBM. We also observe a confounding response to the targeting of CHK and MDM2 activity. CHK kinases are master regulators of the response to DNA damage and the p53 transcription factor to enact cell cycle arrest or apoptosis cell fates. MDM2 is a negative regulator of p53 protein levels, and its inhibition stabilizes p53 levels in the cell. Despite CHK and MDM2 inhibition having opposing effects on p53 activity, both exposures led to a block in the induction of trametinib-induced compensatory transcription. However, p53 is known to negatively regulate CHK1 expression ^76^; therefore, our results may be explained by both exposures leading to a decrease in CHK activity in the cell.

The work presented here constitutes a new approach to prioritize combinatorial therapies based on their induction of gene expression programs of interest. Approaches like sci-Plex-GxE have the potential to complement existing drug discovery pipelines by prioritizing anti-tumor therapies that not only lead to desired anti-proliferative or pro-apoptotic effects but also minimize the possibility of therapeutic resistance. Scrutiny of transcriptional adaptation and its context-dependent genetic requirements could also reveal modules of important genes (e.g. immune checkpoint machinery) that are not sensitive to monotherapy and therefore constitute an orthogonal strategy for treatment. Finally, sci-Plex-GxE reveals the extent to which different modules of genes are coupled through regulation by overlapping signaling pathways, which may shed light on why cells’ responses to drugs or other environmental factors can vary so dramatically.

## Acknowledgments

The authors would like to thank all of the members of the Trapnell and Shendure labs, N.M. Cruz and J.R. McFaline-Figueroa for helpful suggestions, critical discussions, and editing of this manuscript. J.L.M.-F. would like to thank S.V. McFaline-Cruz for their support. The authors would like to thank Lena Christiansen, Fan Zhang, and Frank Steemers for providing reagents and performing sequencing.

## Funding

This work was funded by grants from the NIH (DP1HG007811 and R01HG006283 to J.S.; DP2HD088158 to C.T.), the W. M. Keck Foundation (to C.T. and J.S.), the NSF (DGE-1258485 to S.R.S.), and the Paul G. Allen Frontiers Group (to J.S. and C.T.). J.S. is an Investigator of the Howard Hughes Medical Institute. J.L.M.-F. was previously supported by NIH grants T32HL007828 and T32HG000035. J.L.M.-F is supported by grants from the NIH (R35HG011941) and the NSF (2146007). **Author contributions**: J.L.M.-F., and C.T. conceived the project; J.L.M.-F., S.R.S., and A.J.H. designed experiments; J.L.M.-F., S.R.S, D.J., L.S. performed experiments; J.L.M.-F., S.R.S., and C.T. analyzed the data; J.L.M.-F., C.T., and J.S. supervised the project; and J.L.M.-F., and C.T. wrote the manuscript with input from all authors.

## Data and materials availability

Processed and raw data are available for download from the National Center for Biotechnology Information (NCBI) Gene expression omnibus (GEO) under series number GSE225775. Code used to reproduce the presented analyses is available on github at https://github.com/cole-trapnell-lab/sci-Plex-GxE.

## Declaration of interests

One or more embodiments of one or more patents and patent applications filed by the University of Washington may encompass the methods, reagents, and data disclosed in this manuscript.

## Inclusion and diversity

Three of the authors in this study, including the first author, self-identify as members of groups underrepresented in science. All authors support an equitable, diverse and inclusive conduct of research.

## Methods

### Cell lines, cell culture, and expression of CRISPRi/a systems

A172, T98G, and U87MG glioblastoma cell lines were purchased from ATCC. Cells were cultured in DMEM media (ThermoScientific) supplemented with 10% fetal bovine serum and 1% penicillin/streptomycin (P/S, ThermoScientific). GBM4, GBM8, GSC0131, and GSC0827 glioma stem cell (GSC) cultures have been previously described ^53,77,78^ and were provided by Drs. Robert Rostomily and Andrei Mikheev, University of Washington and Houston Methodist Hospital (GBM4 and GBM8) and Dr. Patrick Paddison, Fred Hutchinson Cancer Research Center (GSC0131 and GSC0827). GSC cultures were maintained in a defined serum-free medium at 37C and 5% O_2_ to mimic *in vivo* conditions. GBM4 and GBM8 were cultured in Neurobasal medium (ThermoScientific) supplemented with B-27 and N2 (ThermoScientific), 20 ng/mL EGF (PeproTech), 20 ng/mL FGF (PeproTech) and 5 µg/mL heparin (Sigma). GSC0131 and GSC0827 were cultured in Neurocult medium (StemCell Technologies) supplemented with 20 ng/mL EGF (PeproTech), 20 ng/mL FGF (PeproTech), and 0.8 µg/mL heparin (Sigma). All cultures were negative for Mycoplasma contamination.

For the generation of CRISPRi-mediated knockdown cells, lentiviral particles encoding dCas9-BFP-KRAB were generated by transfecting HEK293T cells with plasmids encoding dCas9-BFP-KRAB (pHR-SFFV-dCas9-BFP-KRAB, Addgene 46911) and the ViraPower lentiviral packaging mix (ThermoScientific). Transfection was performed using lipofectamine 2000 (ThermoScientific) in OptiMEM (ThermoScientific) following the forward transfection protocol provided by the manufacturer scaled up to 15 cm dishes. 72 hours post-transfection, media was collected and filtered using a 50 mL 0.22 µm steriflip filtration system. A172, T98G, and U87MG cells were then transduced by culturing for 48 hours with different amounts of the filtered lentiviral supernatant. Cells were then expanded, analyzed and sorted using fluorescent activated cell sorting (FACS) for cells with the highest amount of BFP fluorescence starting from transductions with an MOI ∼0.3. To arrive at pure populations of cells with similar levels of dCas9-KRAB cells were expanded and sorted 4 times. For the generation of CRISPRa-mediated overexpression cells we used the 2-component dCas9-SunTag system [citation], a filtered lentiviral supernatant carrying a payload of dCas9-GCN4-BFP (pHRdSV40-dCas9-10xGCN4_v4-P2A-BFP, Addgene 60903) or scFV-GCN4-GFP-VP64 (pHRdSV40-scFv-GCN4-sfGFP-VP64-GB1-NLS, Addgene 60904) were generated as described above. Glioblastoma cells were simultaneously transduced with dCas9-GCN4-BFP at an MOI ∼0.3 and with scFV-GCN4-GFP-VP64 at an MOI ∼1. Cells were expanded, and FACS sorted a total of 4 times based on BFP and GFP fluorescence to ensure maximal and similar expression across cells.

### Generation of CROP-seq-OPTI gRNA libraries

Protospacer sequences targeting all of the perturbed genes in this study were obtained from the genome-wide human CRISPRi and CRISPRa version 2 libraries designed by Horlbeck and colleagues^25^. Oligonucleotides containing these sequences and flanked with adapters with homology to a CROP-seq vector that we have previously altered^23^ to contain a CRIPSRi optimized guide RNA backbone^79^ (CROP-seq-OPTI, Addgene 106280) were synthesized individually for experiments related to **Fig. 1-2** (IDT) and pooled or as a pooled oligo array (CustomArray Inc. Bothell, WA) for our kinome screen.

5‘ homology sequence:

5’-ATCTTGTGGAAAGGACGAAACACC-3’

3’ homology sequence:

5’-GGGTTTAAGAGCTATGCTGGAAACAGCATAGCAAGT-3’

Prior to Gibson assembly, pooled oligonucleotides were amplified via PCR using NEBNext 2X Hi-Fi PCR Master Mix (NEB) and primers:

Forward primer:

5’-ATCTTGTGGAAAGGACGAAACACCG’3’

Reverse primer:

5’-GCTATGCTGTTTCCAGCATAGCTCTTAAAC-3’

Amplification was followed on a MiniOpticon real-time PCR system (BioRad) with the addition of SYBR green (Invitrogen), and reactions stopped prior to saturation. Amplified oligonucleotides were purified using the NucleoSpin PCR clean-up and gel extraction kit (Takara Bio). CROP-seq-opti was linearized via digestion with BsmBI and alkaline phosphatase (NEB) with PCR clean up in between both digestions, purified via gel extraction from a 1% agarose gel followed by cleanup using the NucleoSpin PCR clean up and gel extraction kit (Takara Bio). Linearized CROP-seq-optiI and amplified oligonucleotides were assembled using the NEBuilder HiFi DNA assembly cloning kit (NEB) with the inserts at 2 fold molar excess followed by multiple transformations into NEB stable competent E.Coli (NEB) to ensure at least 20x coverage of colonies for every sgRNA, transformations combined and cultured in 50 mL of Luria broth containing ampicillin at 30°C for 24 hours. Plasmid libraries were recovered using a Midi prep kit (Qiagen). Lentiviral libraries were generated in HEK293T by transfection of plasmid libraries using lipofectamine 2000 (ThermoScientific) in OptiMEM (ThermoScientific) following the forward transfection protocol provided by the manufacturer scaled up to 15 cm dishes. 72 hours post-transfection, media was collected and filtered using a 50 mL 0.22 µm steriflip filtration system. Viral supernatant was titered for each cell line (A172, T98G, and U87MG) by transduction with varying amounts of lentiviral supernatant for 72 hours in 6-well plates. After this, cells were split 1:4 into media with and without 1 µg/mL of puromycin, cultured for 96 hours, and the approximate MOI calculated. For our screens, 3 x 10^6^ cells in 10 cm tissue culture dishes were transduced with lentiviral libraries at an approximate MOI of 0.1 to ensure single integrations. 72 hours post-transduction cells were transferred to two 15 cm tissue culture dishes containing 1 µg/mL of puromycin and continuously cultured in puromycin. Cells were seeded for chemical exposure between 10 to 14 days after transduction.

### Chemical exposure of genetically perturbed cell pools

Temozolomide (cat no. T2577) and 6-thioguanine (cat no. A4882) were purchased from Sigma and resuspended in DMSO (VWR scientific) to a concentration of 100 mM. Lapatinib (cat no. S2111), nintedanib (cat no. S1010), trametinib (cat no. S2673), and zstk474 (cat no. S1072) were purchased from Selleck Chemicals at a concentration of 10 mM in DMSO. Genetically perturbed pools of glioblastoma cells were seeded in 96-well plates at 2.5 x10^4^ cells per well in 100 µL of DMEM containing 10% FBS, 1% P/S, and 1 µg/mL puromycin and allowed to attach for 24 hours. Small molecules were diluted to 1000-fold the exposure concentration in DMSO, followed by a 10-fold dilution into Dubelcco’s Phosphate buffered saline (DPBS, Life Technologies) and 1 µL added of the appropriate drug and dose to wells of seeded cells and a final concentration of 0.1% v/v DMSO. For temozolomide and 6-thioguanine exposure experiments, cells were exposed for 96 hours. For lapatinib, nintedanib, trametinib and zstk474 experiments, cells were exposed for 72 hours.

### sci-Plex cell harvest and hash labeling

Cell harvest and sci-Plex labeling were performed as previously described^13^. Briefly, drug-containing media was removed from wells, wells were washed with 100 µL of DBPS, and 50 µL of TrypLE (Invitrogen) was added to every well. Cells were detached by incubation with TrypLE at 37°C. Once cells were detached, 100 µL of ice-cold DMEM was added to every well, cells resuspended, cells transferred to v-bottom 96 well plates, cells pelleted by centrifugation and washed with ice-cold DPBS. Cells were lysed to nuclei by the addition of 50 µL of cold lysis buffer (CLB: 10 mM Tris HCl ph 7.4, 10 mM NaCl, 3 mM MgCl2, 0.1% IGE-PAL) containing 1% v/v Superase-In and 100 nM (final concentration) hashing oligos (each unique to each well) of the form:

5’-GTCTCGTGGGCTCGGAGATGTGTATAAGAGACAG-[10bp-barcode]-BAAAAAAAAAAAAAAAAAAAAAAAAAAAAAAAA-3’

Where B is G, C or T (IDT), lysis was carried out on ice for 3 minutes, followed by the addition of 200 µM of 5% paraformaldehyde (EM solutions) in 1.25x PBS and incubated on ice for 15 minutes. Nuclei were then pooled, pelleted by centrifugation, and washed twice with 2 mL of CLB containing Superase-In and 1% v/v of 20 mg/mL molecular grade BSA (NEB). After the final wash, nuclei were resuspended in 1 mL of CLB containing Superase-In and 1% v/v of 20 mg/mL molecular grade BSA and snap-frozen in liquid nitrogen. Labeled nuclei were stored at –80°C until the preparation of sequencing libraries.

### Preparation and sequencing of single-cell RNA-seq and CROP-seq-OPTI sgRNA enrichment libraries

Flash-frozen nuclei were thawed at room temperature, nuclei pelleted by centrifugation at 500 x *g* for 5 minutes, the supernatant removed, nuclei re-suspended in 1 mL of CLB containing 1% v/v Superase-In and 1% v/v of 20 mg/mL molecular grade BSA (NSB) and nuclei from uniquely hashed samples were pooled. Pooled nuclei were then pelleted by centrifugation at 500 x *g* for 5 minutes. For a subset of experiments, the same hashes were used for different replicates and/or cell lines. As such, these were not combined and distributed across unique wells of the plate in which reverse transcription (RT) was performed (e.g. for cells exposed to inhibitors and damaging agents that alter cell stress pathways each line was hashed separately and each cell line arrayed across 4 columns of the 96-well RT plate). Prior to RT, nuclei were further permeabilized by incubation in 0.2% tryton-X100 (Sigma) in NSB. Nuclei were pelleted, resuspended in 400 µL of NSB, and sonicated for 12 seconds using the low setting on a Bioruptor sonicator (Diagenode). Nuclei were then pelleted, resuspended in 500 µL NSB, stained with trypan blue (Life Technologies), and counted on a hemocytometer. Nuclei distributed into skirted lo-bind 96 well plates (Eppendorf) at 20,000 (related to figures 1 & 2) or 40,000 nuclei per well in 22 µL of NSB and 2 µL of 10 mM dNTP mix (NEB).

To increase our rate of sgRNA assignment, we devised a sgRNA enrichment strategy specific to combinatorial indexing sci-RNA-seq that relies on (1) the addition of a custom RT primer targeting the sgRNA-containing puromycin transcript delivered by CROP-seq, (2) performing combinatorial indexing solely on the i5 end of the mRNA molecule, and (3) addition of a sgRNA enrichment PCR from the final mRNA library which targets the sgRNA-containing puromycin transcript while maintaining the combinatorial i5 cell barcode on every molecule (**Fig. 1A**). We designed the targeted RT primer to capture transcripts derived from CROP-seq-OPTI ^23^, a modified version of CROP-seq incorporating an optimized single-guide RNA backbone ^79^ that increases the stability of sgRNA association with dCas9.

For our sci-Plex-GxE protocol, RT was performed as previously described^13^ with the addition of 2 µL of 100 µM ligation-compatible indexed oligo-dT primer of the form:

5′-/5Phos/CAGAGCNNNNNNNN-[10bp-barcode]-TTTTTTTTTTTTTTTTTTTTTTTTTTTTTT-3′,

Where N is any base (IDT) and 1 µL of 100 µM ligation compatible indexed CROP-seq-OPTI targeting primer of the form

5’-/5Phos/CAGAGCNNNNNNNN-[10bp-barcode]-ACTTTTTCAAGTTGATAACGGACTAGCCTT ATTT-3’

Where N is any base (IDT) that were added to every well. The use of the OPTI-modified backbone necessitated additional considerations for the design of the targeted RT primer to ensure that the hairpin that mediates strong binding to dCas9 does not interfere with the efficiency of reverse transcription. Primers were annealed by incubation at 55°C for 5 minutes, followed quickly by incubation on ice. 14 µL of RT mix (8 µL of Superscript IV buffer, 2 µL of Superscript IV enzyme, 2 µL of 100 mM DTT and 2 µL of RNAseOut rnase inhibitor, Invitrogen) were added to each well, and RT performed as follows: 4°C - 2 min, 10°C - 2 min, 20°C for 2 minutes, 30°C for 2 minutes, 40°C for 2 minutes, 50°C for 2 minutes and 55°C for 15 minutes. After RT, 60 µL of CLB containing 1% v/v of 20 mg/mL molecular grade BSA (NBB) were added to every well, wells pooled, nuclei pelleted, resuspended in NSB and 10 µL of nuclei were redistributed into each well of a 96 well Lo-bind skirted plates. All experiments were done using a single RT and ligation plate with the exception of the kinome screen where 4 RT and 4 ligation plates were used. For the second round of combinatorial indexing, 8 µL of indexed ligation primer of the form

5’-GCTCTG[9bp-or-10bp-barcode-A]/ideoxyU/ACGACGCTCTTCCGATCT[reverse-complement- of barcode-A]-3’

(IDT) were added to each well, followed by the addition of 22 µL of ligation mix (20 µL quick ligase buffer and 2 µL of quick ligase, NEB) and incubation at 25°C for 10 minutes. After ligation 60 µL of NBB were added to each well, wells pooled, nuclei pelleted by centrifugation at 700 x *g* for 10 minutes, washed twice with NBB, nuclei counted, and redistributed into 96 well Lo-bind skirted plates. The number of cells distributed was determined by the number of RT and ligation barcodes in the experiment so as to minimize the number of total doublets in the experiment to between 1-10% and the rate of doublets that cannot be filtered from sci-Plex hashes to 1% or less according to birthday problem statistics^80^. Plates were stored at –80°C until further processing. Second strand synthesis was performed after thawing by the addition of 5 µL of second strand synthesis mix (3 µL of elution buffer [Qiagen], 1.33 µL mRNA second strand synthesis buffer and 0.66 µL of second strand synthesis enzyme mix [NEB]) and incubated at 16°C for 3 hours. After second strand synthesis, DNA was tagmented by the addition of 10 µL of tagmentation mix (0.01 µL of a custom TDE1 enzyme in 9.99 µL of 2x Nexterda TD buffer, Illumina) and plates incubated at 55°C for 5 minutes. After tagmentation, 20 µL of DNA binding buffer (Zymo) was added to each well and incubated at room temperature for 20 minutes, 40 µL of Ampure XP beads (Beckman Coulter) was added to every well, and a cleanup was performed according to manufacturer’s instructions with changes to the elution step. Prior to elution, beads were incubated with 10 µL of USER mix (1 µL of 10X USER buffer and 1 µL of USER enzyme in 8 µL of nuclease-free water, NEB) and incubated at 37°C for 15 minutes. After incubation, 7 µL of elution buffer was added to each well, beads were resuspended, plates were placed on a magnetic stand and 16 µL of solution was transferred to 96 well Lo-bind skirted plates. For PCR, 20 µL of 2X NEBNext master mix, 2 µL of 10 µM indexed P5 primer of the form:

5′-AATGATACGGCGACCACCGAGATCTACAC-[index5-]ACACTCTTTCCCTACACGACGCTCT TCCGATCT-3′

and 2 µL of 10 µM indexed P7 primer of the form:

5′-CAAGCAGAAGACGGCATACGAGAT-[index7]-GTCTCGTGGGCTCGG-3′

were added to each well. To account for the loss of the P7 index during sgRNA enrichment PCR, each PCR plate was labeled with 96 unique P5 indices, and the P7 index was used as a plate identifier. Libraries were generated using the following PCR program: 72°C for 5 min, 98°C for 30 sec, 15 cycles of (98°C for 10 sec, 66°C for 30 sec, 72°C for 30 sec), and a final extension at 72C for 5 minutes. After PCR, uniquely labeled wells were pooled, and 1 mL of PCR product was subjected to a 0.7X Ampure cleanup. After the initial incubation, the supernatant was transferred to a new tube, and additional beads were added to arrive at a 1X Ampure cleanup which will be the hash-containing fraction. Both fractions were further processed following the standard Ampure XP protocol and eluted in 100 µL of elution buffer.

For the enrichment of sgRNA containing library fragments, a separate sgRNA enrichment PCR was performed via nested PCR using the final sci-RNA-seq3 libraries as starting material. For each library, 10-20 unique reactions were performed each using 1:100^th^ of the mRNA library in a reaction containing 25 µL of 2X NEBNext master mix an up-stream U6 targeting forward primer of the form 5′- CTTGTGGAAAGGACGAAACACCG-3′, a reverse primer targeting the P5 flow cell binding sequence (5′- AATGATACGGCGACCACCGA-3′), 0.5 µL of SYBR green (Life Technologies) and nuclease-free water. Amplification was monitored by real-time PCR (BioRad), PCR terminated during the extension phase just prior to saturation, PCR was purified using a 1X Ampure XP cleanup, and eluted into 50 µL. A second PCR reaction was performed as described above with the following forward primer targeting the sgRNA proximal U6 promoter and containing an Illumina read 2 primers binding sequence (5′- GTCTCGTGGGCTCGGAGATGTGTATAAGAGACAGCTTGTGGAAAGGACGAAACACCG-3′) and reverse primer targeting the P5 flow cell binding sequence followed by a 1X Ampure cleanup. Finally a third PCR was performed using a P7 index as above that could be used to link an mRNA library to its corresponding sgRNA enrichment library and the reverse primer targeting the P5 flow cell binding sequence followed by a 1X Ampure cleanup.

Library fragment sizes were determined using an Agilent TapeStation high sensitivity screen tape (Agilent) and library concentration determined using a Qubit fluorometer (Life Technologies). Libraries were sequenced on the NextSeq 550 (R1: 34 bp, R2: 100 bp, I1: 10 bp, I2: 10 bp), Nextseq 2000 (R1: 34 bp, R2: 70 bp, I1: 10 bp, I2: 10 bp) and Novaseq (R1: 34 bp, R2: 100 bp, I1: 10 bp, I2: 10 bp) platforms.

### Data processing and generation of count data matrix

Sequences were demultiplexed using bcl2fastq (Illumina) filtering for reads with RT and ligation barcodes within an edit distance of 2 bp. PolyA tails were trimmed using trim-galore (https://github.com/FelixKrueger/TrimGalore) and reads were mapped to the human hg-38 transcriptome using STAR^81^. After alignment, reads were filtered by alignment quality and duplicates were removed if they mapped to the same gene, the same barcode and the same unique molecular identifier (UMI) or if they met the first 2 criteria and the UMI was within an edit distance of 1 bp. Reads were assigned to genes using bedtools^82^. 3’ UTRs were extended by 100 bp in the gene model to account for short 3’ UTR annotations to minimize genic reads labeled as intergenic. A knee plot was used to set a threshold above which a combinatorial cell barcode confidently corresponded to a cell. UMI counts for cell barcodes that pass this threshold were aggregated into a sparse matrix format, followed by the creation of a cell data set object using Monocle3. Mitochrondrially encoded genes were excluded in downstream analyses.

### Hash and sgRNA assignment

sci-Plex hashes and sgRNA containing puromycin transcripts derived from CROP-seq were isolated from demultiplexed reads. Hashes were assigned as previously described ^13^. Briefly, reads were considered hashes if (1) the first 10 bp of read 2 matched a hash in a hash whitelist within a hamming distance of 2 and (2) contained a poly A stratched spanning the 12-16 base pair region of read 2. For sgRNA assignment, read were considered CROP-seq derived if the bases spanning position 24-42 matched a sgRNA in a sgRNA whitelist within a hamming distance of 2 and (2) a TGTGG sequence at position 3-7 of read 2. Duplicated reads were collapsed by their UMIs arriving at hash and sgRNA UMI counts for each nucleus in our experiment. Finally, we tested whether a particular nucleus was enriched for one or more particular hash or sgRNA as described in ^13^ for sci-Plex hashes.

### Data pre-processing

For our kinome screen, multiplets were removed from our experiments using 3 orthogonal approaches. First, doublets were inferred using scrublet^83^ specifying an expected doublet rate of 0.05 as calculated using a formulation of the birthday problem. Cells with a doublet score of larger or equal to 0.25 were removed from our dataset ( 0.88% of cells). Next cells where the ratio between the UMI counts of the most abundant and next most abundant hash (i.e., the top to second best ratio) was less than 2.5 or cells with less than 5 totals hash UMIs were removed from our analysis ( 4.9% of cells). Lastly, all of the cells from our experiment were co-embedded in a UMAP[1] projection. Data were pre-processed by performing an initial dimensionality reduction using principal component analysis (PCA) using genes expressed in at least 5% of the cells from each cell line as feature genes and the top 25 dimensions were used to build our UMAP. We specified 20 nearest neighbors and a minimum distance of 0.1 as UMAP hyperparameters. We next clustered cells in this co-embedding using Leiden community detection^84^ specifying a resolution parameter of 1e-6. This resulted in 5 UMAP partitions that could be readily assigned to the 3 cell lines in our experiment by visual inspection. This approach identified a small proportion of cells[2] where there was a mismatch between hash and transcriptome identity. For our GSC and chemical exposure experiments, multiplets were described as above without scrublet pre-filtering. For our proof-of-concept experiments in A172 cells, multiplets were removed using the hash filters described above.

### Differential gene expression analysis

Differential gene expression analysis was performed using the *fit_models* function in Monocle3. For defining the effect of drug exposure on the gene expression profiles of unperturbed cells, we created subsets of our dataset for every exposure and set of NTC cells. For every gene expressed in at least 5% of cells, we fit a generalized linear model of the form expression ∼ log(dose + pseudocount) specifying “∼ log(dose + 0.001)” for the model_formula_str parameter in fit_models. We then combined all tests for all genotypes and doses and corrected for multiple hypothesis testing using the Benjamini-Hochberg false discovery rate method. We chose a pseudocount of 0.001 to preserve the relationship to a dose with minimal effect on cells based on preliminary experiments (data not shown).

For differential gene expression analyses, gene expression was log transformed after addition of a pseudocount of 1. For experiments where we exposed perturbed A172 cells to temozolomide, we created subsets of our dataset for every dose of temozolomide and every pairwise combination of a target and NTC cells. For all expressed genes, we fit a generalized linear model of the form expression ∼ genotype specifying “∼ gene_id” for the model_formula_str parameter in fit_models. We then combined all tests for all genotypes and doses and corrected for multiple hypothesis testing using the Benjamini-Hochberg false discovery rate method. To determine differential kinase expression as a function of kinase perturbation, we created subsets of our data for every pairwise combination of target and NTC cells. We next fit a generalized linear model of the form expression ∼ genotype + log(UMI counts) + replicate specifying “∼ gene_id + replicate + log(n.umi)” for the model_formula_str parameter in fit_models and performed multiple hypothesis testing as above. To determine differential kinase expression as a function of kinase perturbation and drug exposure, we created subsets of our data for every exposure and every pairwise combination of target and NTC cells. We next fit a generalized linear model of the form expression ∼ genotype + log(dose + pseudocount) + replicate specifying “∼ gene_id + replicate + log(dose + 0.001)” for the model_formula_str parameter in fit_models and performed multiple hypothesis testing as above. Aggregation of gene expression counts and scoring cells by levels of expression signatures

### Gene set analysis

Gene set enrichment analysis was performed using the piano R package^85^. The hallmarks and oncogenic signatures^57,86^ gene sets were obtained from the Broad Institute’s Molecular Signatures Database^56^. Hypergeometric testing was performed using feature genes as the foreground and all genes as the background.

### UMAP embedding of knockdown proportions

We used UMAP to visualize the relationship between genotypes and the proportion of temozolomide and perturbation-induced cellular states. We performed an initial dimensionality reduction using PCA returning the top 25 principal components using the union of all differentially expressed genes as a function of genotype as feature genes. Genes that were differentially expressed between negative controls and NTC cells were removed from the analysis. We then performed dimensionality reduction using UMAP, specifying 20 nearest neighbors and a minimum distance of 0.1 as hyper-parameters. We clustered cells within the UMAP embedding using Leiden community detection^84^. We next calculated the frequency of cells for each gRNA or genotype across clusters and used these matrices to initialize a UMAP embedding.

### Calculation of pairwise angular distance

To detect the effect of a genetic perturbation on a drug-induced transcriptional program we calculated the pairwise angular distance of every cell to the average profile of non-targethe ting cells exposed to the highest dose of each drug. The angular distance between two cells was calculated as the arc cosine transcriptome distance between the norm of the expression vector for each cell over a set of feature genes such that if *V* is the expression vector of a cell across *x* gene expression values, then the norm of the vector is defined as,

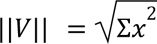

and the angular distance between the vector norms for two cells is,

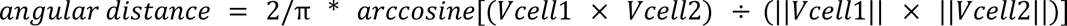

For all angular distance calculations in perturbed cells the comparisons were made between every cell (perturbed and unperturbed) vs. the mean profile of NTC cells exposed to the highest dose of each drug. We compared this approach to the use of the more common Jensen-Shannon distance metric, observing good agreement between both distance metrics (**Supp Fig. 3G**). Therefore, we chose to continue with angular distance, which is a less expensive calculation, for our measure of similarity to unperturbed cells.

### Inference of the relative transcriptional effective concentration 50 (TC50)

First, we fit a 4-parameter log-logistic dose-response model to the relationship between the mean pairwise angular distance and dose of temozolomide for NTC control cells using the drc R package^87^. We specified a formula of mean angular distance ∼ dose and the function as LL.4 in the drm function. We then estimated the effective dose 50, which we term our transcriptional effective concentration 50 (TC50) using the ED function of the drc package. For each genotype, we fit a line to the relationship between the log of the mean angular distance to NTC vs the log of the dose of temozolomide. We then used those fits to determine the concentration at which cells reached the mean angular distance to NTC that NTC cells achieved at their TC50.

### Median kinase knockdown

We assessed the quality of our gRNA assignments in our kinome screen by examining the median knockdown level across all perturbed kinases in our experiment. We calculated the mean expression levels for each kinase in NTC cells and their respective perturbed target cells at varying gRNA read cutoffs (i.e. 1-10 gRNA reads per cell). We ensured that our knockdown estimates were not biased due to the zero inflation of sc-RNA-seq data and the larger proportion of NTC cells in our experiment by permuting the gRNA-defined target labels and re-calculating mean kinase knockdown. We then compared the distribution of knockdown levels between our gRNA assigned and permuted data using the non-parametric two-sample Wilcoxon test.

### Enrichment and depletion of knockdowns

We assessed the relative enrichment and depletion of kinases in our experiment by comparing the frequency of gRNAs targeting a particular kinase in our final dataset to its frequency in the plasmid library used to create gRNA-delivering lentiviral particles. We defined the relative proportion of gRNA against a kinase target as the mean-centered log of the ratio of the two frequencies.

### Identifying conserved responses to RTK pathway inhibition

We determined conservation in response to RTK pathway inhibition by comparing the dynamics of differential gene expression of unperturbed A172, T98G, and U87MG cells. The mean expression in NTC cells of differentially expressed genes as a function of lapatinib, nintedanib, trametinib, and zstk474 exposure was clustered for each cell line individually by hierarchical clustering. We chose k = 6 as the number of clusters for each cell line by visual inspection of dendrograms across all cell lines. We then calculated the Jaccard coefficient for every pairwise comparison of clusters across all 3 cell lines. Clusters with a Jaccard coefficient over 0.1 were collapsed into conserved super-clusters by taking the union of the genes across similar clusters.

### Estimation of the cell cycle stage of single-cells

Estimates for the cell cycle stage of individual cells was inferred as in^13^. Briefly, the expression of genes associated with the G1/S and G2/M cell cycle stages was size-factor normalized ^88^, and their expressions aggregated and log-transformed. We define a proliferation index as the sum of the logged G1/S and G2/M scores.

### Chemical genomic validation of kinases whose loss leads to changes in the induction of the compensatory adaptive program

To validate the contribution of kinase hits to the induction of the compensatory adaptive program, we exposed A172, T98G and U87MG to one of 23 compounds in the absence or presence of trametinib, the strongest inducer of the adaptive compensatory program. Cells were exposed to 0.01, 0.1, 1, and 10 µM doses of each compound, the absence or presence of 0.01, 0.1, 1 and 10 µM of trametinib or DMSO vehicle control. For trametinib co-exposure conditions, the concentrations were matched for each compound and trametinib (e.g, 1 µM of compound + 1 µM trametinib). The concentration of DMSO control was set to 0.2% v/v across all single and combinatorial exposures. Cells were exposed to compounds for 72 hours, harvested, multiplexed using our previous sci-Plex protocol ^13^, and nuclear mRNA libraries were generated and sequenced as described above.

### Correlation of single and combinatorial chemical targeting to the effect induced by trametinib exposure

For our validation chemical genomic experiment, we performed differential expression analysis using quasipoisson regression for the effect of each compound alone or in combination with trametinib on the set of genes that compose the compensatory adaptive program. For each cell line, we fit a generalized linear model of the form *expression ∼ log(dose + 0.001) + replicate*.

For single chemical exposures, we focused on treatments that led to significant changes in the expression of at least 100 genes of the program (FDR < 5% and a normalized beta coefficient for the dose term of |β_coef_| > 0.05) across two or more cell lines. We calculated the pairwise Pearson’s correlation across all exposures on a matrix of the normalized β coefficients for the dose term across all feature genes. We defined compound exposures with a significant correlation to the effect induced by trametinib exposure as those with a Pearson’s ⍴ > ± 0.2 at an FDR < 0.05.

For both single and combinatorial exposures, we also broadly examined the correlation structure across exposures. We regressed the effect of cell line background for each correlation matrix using the monocle3 function *align_cds* specifying a *residual_model_formula_str* of “cell_line”. We used Leiden-based community detection as implemented in the *cluster_cells* function of monocle3 on this corrected correlation matrix to identify groups of exposures that lead to similar transcriptional changes across genes that make up the compensatory adaptive transcriptional program. To visualize our results, we used this corrected correlation matrix to initialize a UMAP embedding using the *reduce_dimension* function of monocle3 specifying *umap.n_neighbors* of 5 and a *umap.min_dist* of 0.15.

## Supplementary Figures

**Supplementary Figure 1.**
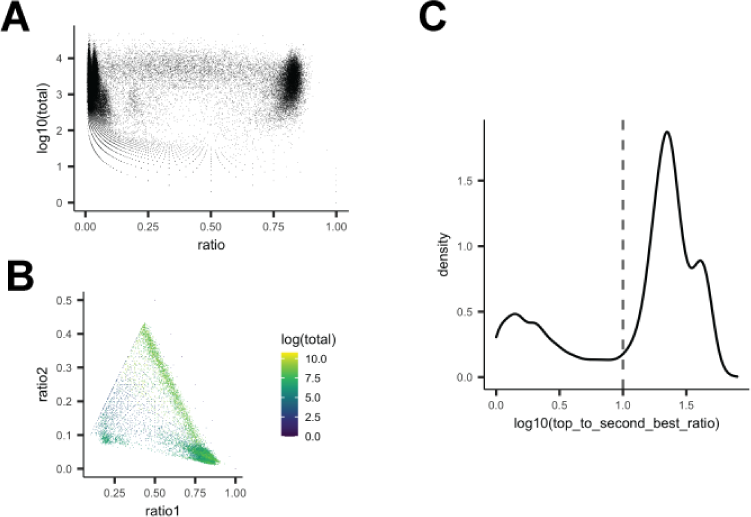
Sensitivity and specificity of sgRNA capture in the context of combinatorial indexing RNA-seq with sci-Plex-GxE. (**A**) Plot of the log10 of the total number of sgRNA reads for a given sgRNA in a given cell as a function of the ratio of that sgRNA to all other sgRNAs in a cell (ratio). (**B**) Plot of the relationship between the proportions of the sgRNA with the highest number of reads in a cell (ratio1) vs the second most prevalent sgRNA (ratio2). (**C**) Density plot of the log10 of the top to second best ratio, the ratio between the proportion of the top most prevalent sgRNA to the second most prevalent.

**Supplementary Figure 2.**
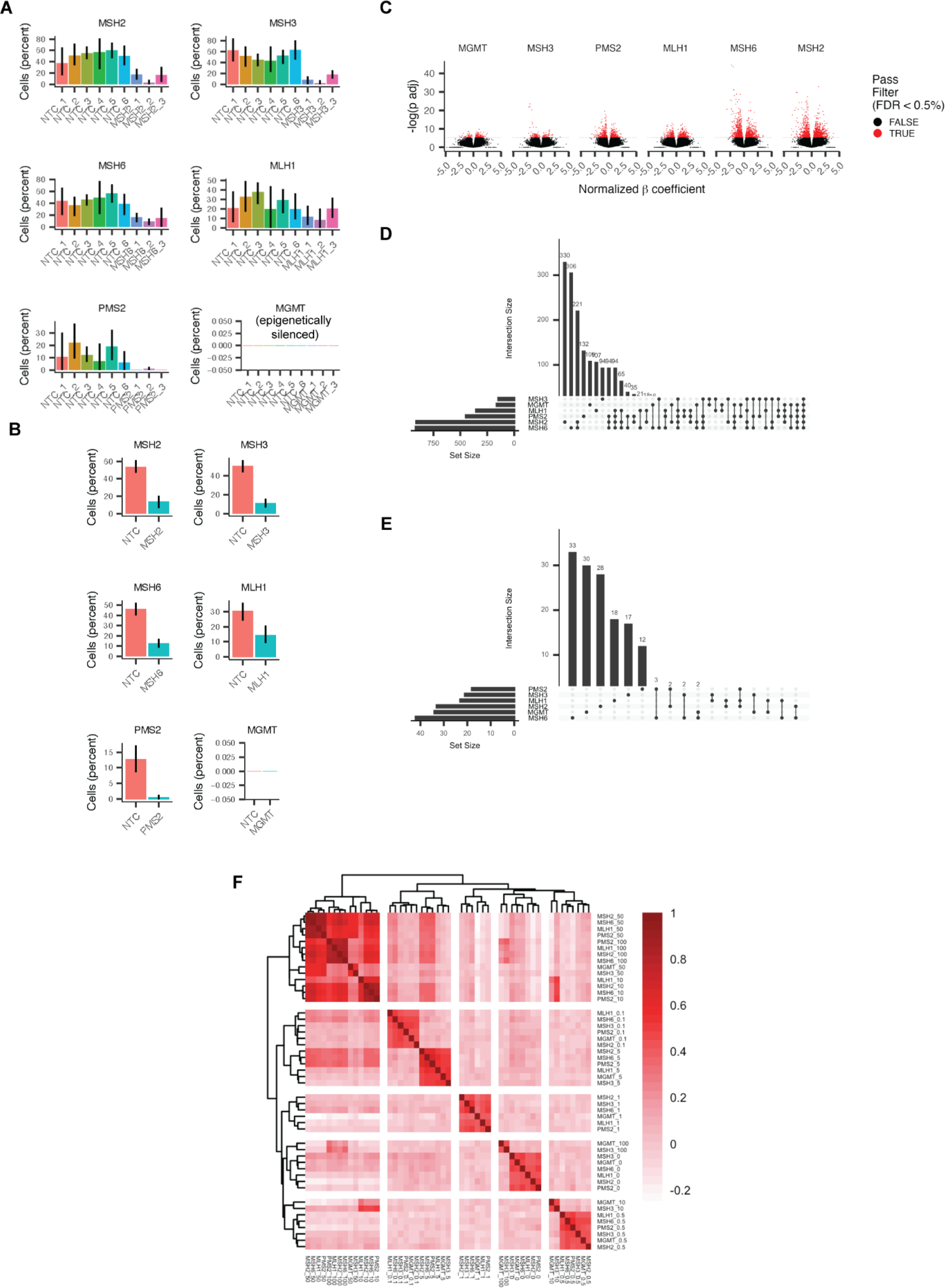
Effects of mismatch repair perturbation of gene expression changes induced by exposure of glioblastoma cells to the chemotherapeutic agent temozolomide. (**A**) Bar plots of the expression level of the mismatch repair components *MSH2*, *MSH3*, *MSH6*, *MLH1*, *PMS2* and the direct repair enzyme MGMT in A172 dCas9-KRAB cells expressing individual sgRNAs against each gene or non targeting controls sgRNAs (NTC). Note that MGMT is not expressed in A172 cells due to epigenetically silencing of the *MGMT* locus by promoter methylation (citation). (**B**) Bar plots as in (**A**) across cells binned by their respective target. (**C**) Volcano plots of the relationship between statistical significance and effect size for the results of differential gene expression analysis of the effect of genetic perturbation of *MSH2*, *MSH3*, *MSH6*, *MLH1*, *PMS2,* and *MGMT* on gene expression after exposure to various doses of temozolomide. For each genotype, genetically perturbed and NTC cells were subsetted by dose, expression was log-transformed and differentially expressed genes were identified by fitting a generalized linear model of the form *expression ∼ genotype*. All tests were then combined and p-values corrected for multiple hypothesis testing using Benjamini-Hochberg. Red indicates genes whose temozolomide-induced gene expression is significantly affected by genetic perturbation at FDR < 0.05. (**D-E**) Upset plots of the overlap of differentially expressed genes in the presence (**D**) or absence (**E**) of temozolomide exposure. (**F**) Correlation heatmap of the normalized effect sizes across all perturbation-dependent differentially expressed genes upon exposure to temozolomide. Rows and columns are labeled as “genotype_dose of temozolomide”. Pearson’s correlation.

**Supplementary Figure 3.**
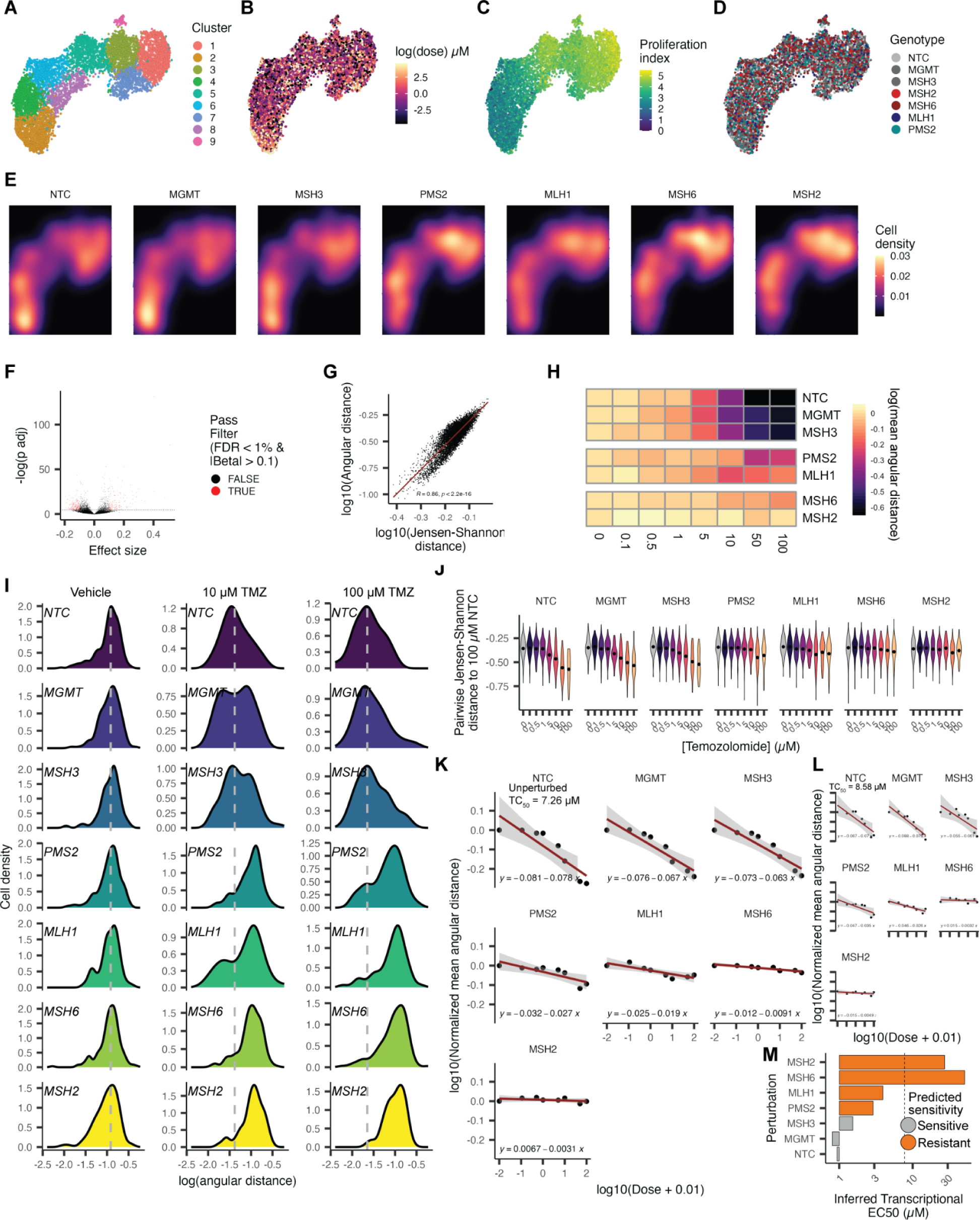
Summarizing state-level effects of temozolomide exposure as a function of genetic perturbation. **A-D**) UMAP embedding of vehicle and temozolomide exposed and genetically perturbed A172 dCas9-KRAB cells. Cells are colored as a function of PCA cluster (**A**), dose of temozolomide (**B**), aggregate expression of genes associated with proliferation (**C,** proliferation index), or genotype (**D**). **E**) Density of cells across the UMAP embedding from (**A-D**). **F**) Volcano plot the relationship between statistical significance and effect size for the results of differential gene expression analysis of the exposure of NTC cells to temozolomide using a model of expression ∼ log(dose of temozolomide + 0.01). **G**) Correlation of the pairwise distances calculated using cosine angular distance or Jensen-Shannon distance between every cell and the mean expression of NTC cells exposed to 100 µM TMZ. **H**) Heatmap depicting the log of the mean angular distance between between every cell and the mean expression of NTC cells exposed to 100 µM TMZ for every genotype dose combination. **I**) Distribution of pairwise angular distance to the mean expression of NTC cells exposed to 100 µM TMZ for genetically perturbed cells exposed to vehicle (left panel) or exposed to 10 µM (middle panel) or 100 µM (right panel) temozolomide. Grey vertical lines refer to the median angular distance of unperturbed NTC control cells for each dose of temozolomide. **J**) Violin plots of the pairwise Jensen-Shannon distance between every cell and the mean expression of NTC cells exposed to 100 µM TMZ as a function of dose of temozolomide to which cells for every genotype were exposed to. **K**) Linear regression fits between dose and mean pairwise angular distance for all genotypes in our experiment. EC50 denotes the concentration at which we observed half of the shift in pairwise angular distance of temozolomide exposed NTC cells obtained from a four parameter log-logistic regression. **L**) Linear regression fits as in K after subsampling genetically perturbed cells (*MSH2*, *MSH3*, *MSH6*, *MLH1*, *PMS2,* and *MGMT* perturbed) to 25 cells per genotype. Panels are ordered as in **K**. **M**) Inferred transcriptional effective concentration (EC50) defined as the concentration of drug necessary to reach 50% of the change in angular distance exhibited by TMZ-exposed NTC cells after subsampling from L. Dashed line: maximum molarity of an aqueous solution as a threshold for genotypes where drug cannot induce 50% of the effect observed in NTC.

**Supplementary Figure 4.**
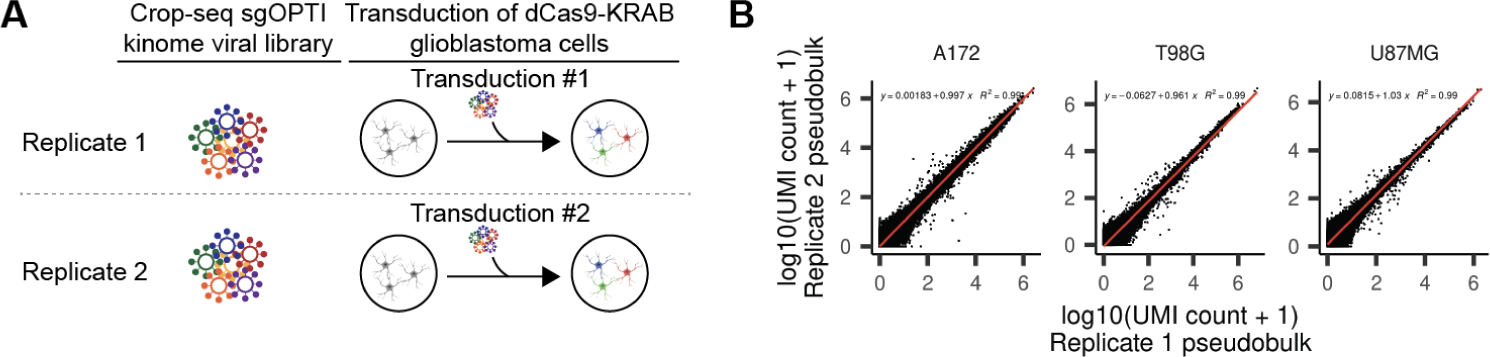
Experimental design and QC. **A**) Schematic depicting the replicate structure of our single-cell kinome screen. **B**) Correlation between replicate screens across the three glioblastoma cell lines in our experiment.

**Supplementary Figure 5:**
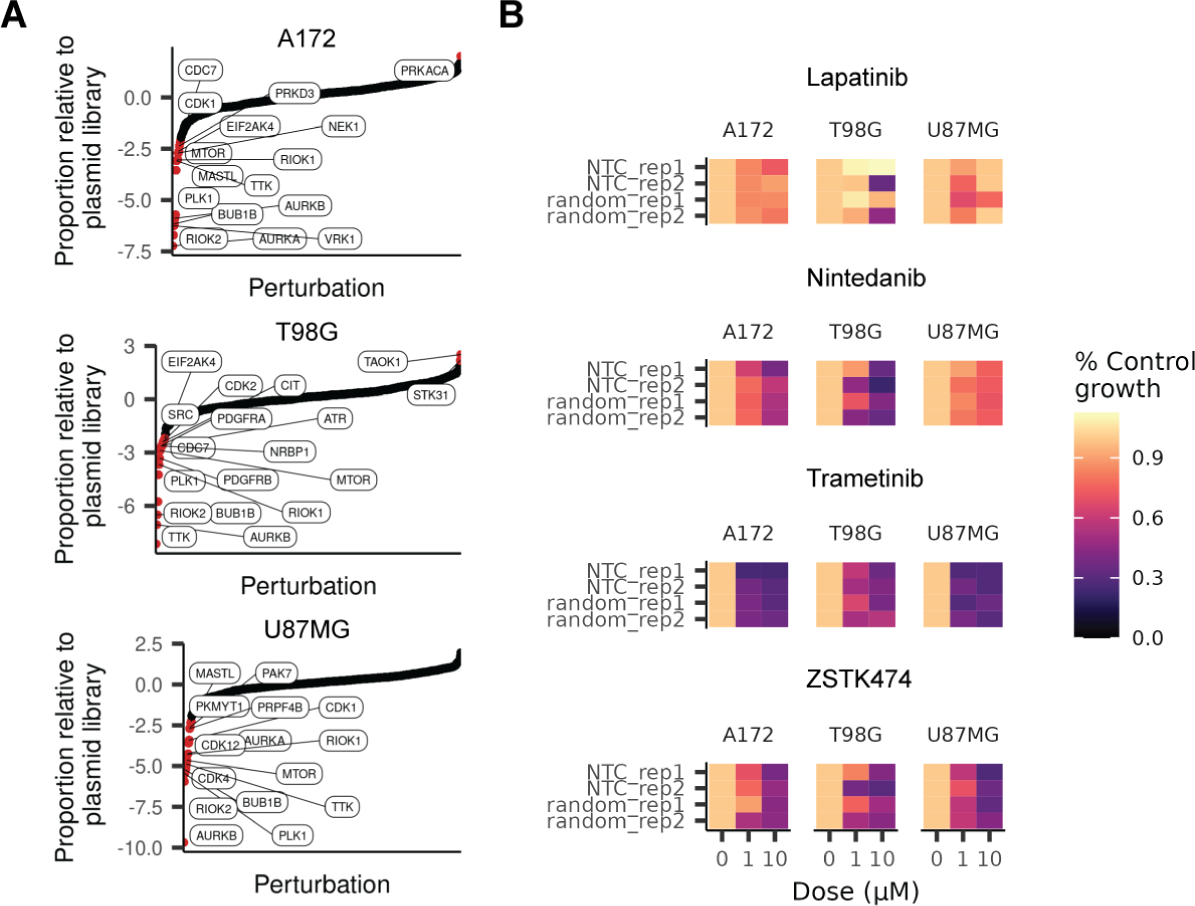
Effect of kinase perturbation and RTK pathway inhibition on GBM cell bulk viability. A. Proportion of cells expressing sgRNAs targeting individual kinases in our screen relative to the starting proportion of sgRNAs in our CROP-seq kinome plasmid library. Labels correspond to depleted kinases over a z-score of 1. **B.** Heatmap depicting viability estimates derived from sci-Plex cell counts for unperturbed cells expressing non-targeting or random targeting control sgRNAs.

**Supplementary Figure 6.**
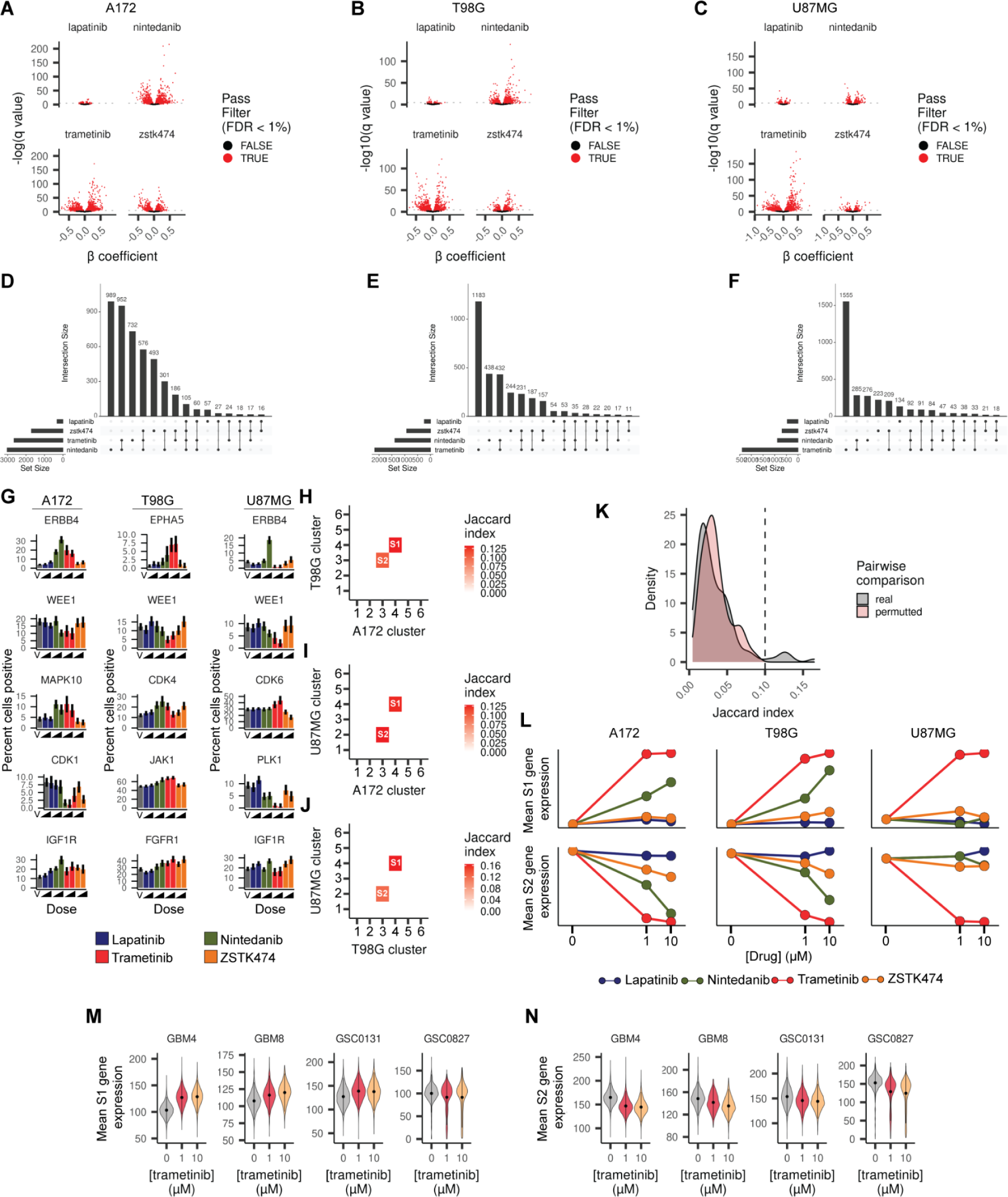
Exposure to small molecule inhibitors targeting the RTK pathway leads to dynamic rewiring of transcriptional networks in glioblastoma cells. **A-C**) Volcano plots of the relationship between statistical significance and effect size of exposure to small molecules targeting the RTK pathway (lapatinib, nintedanib, trametinib, zstk474) on gene expression for A172 (**A**), T98G (**B**), and U87MG (**C**) cells. A generalized linear model of the form expression ∼ log(dose) + replicate was fitted across log normalized expressions for every subset containing cells exposed to one of the 4 agents and vehicle control. Red dots correspond to genes whose expression is significantly altered by kinase perturbation at an FDR < 0.01. D-F) UpsetR plots depicting the overlap in differentially expressed genes as a function of exposure to RTK pathway targeting agents in A172 (**D**), T98G (**E**), and U87MG (**F**) GBM cells. **G**) Bar plots depicting the percent of cells positive for the specified kinase transcripts differentially expressed in at least 1 of 4 drug exposures across the 3 GBM lines in our study. **H-J**) Heatmaps depicting the top overlap of genes between cell clusters for each pair-wise combination of teh 3 GBM cell lines. Significant overlap is defined as having a Jaccard coefficient larger than 0.1 (details in **K**). **K**) Distribution of Jaccard indices for the overlap between drug-responsive gene clusters from Fig. 4A-**C**. Real vs. permutted refers to the Jaccard indices calculated between clusters without and with perturbation of gene ids. Based on this test, we used a Jaccard index of 0.1 or larger to collapse signature genes across cell types. **L**) Aggregate expression of conserved upregulated (S1) and downregulated (S2) gene signatures as a function of each dose, exposure, and cell type. **M-N**) Violin plots depicting the expression of conserved S1 (**M**) and S2 (**N**) signatures in trametinib exposed glioma stem cell cultures.

**Supplementary Figure 7.**
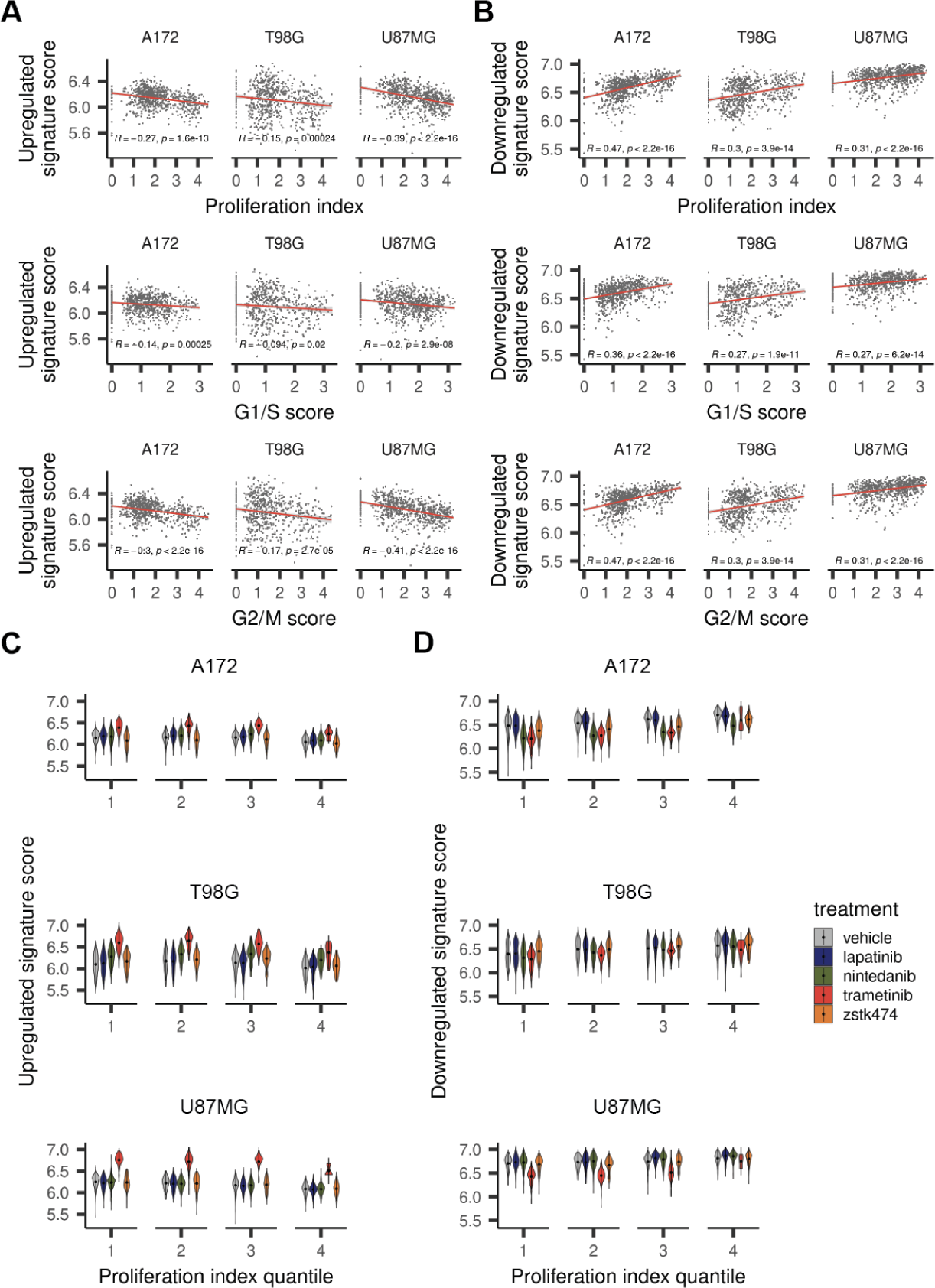
Conserved drug-dependent signatures are correlated with the cell cycle stage of a cell and by MEK kinase activity across all stages of the cell cycle. **A-B**) Expression of conserved S1 (**A**) and S2 (**B**) signatures across NTC cells as a function of the aggregate expression of genes associated with proliferation (top panels), G1/S (middle panels), and G2/M (lower panels) phases of the cell cycle. **C**) Violin plots depicting the expression of S1 (**C**) and S2 (**D**) signatures as a function of treatment across cells of varying proliferation index quantiles (i.e. the x-axes of the top panel of **A** and **B** were divided into 4 equal-sized bins). Note that trametinib exposed cells have higher S1 expression and lower S2 expression across all bins compared to vehicle control.

**Supplementary Figure 8.**
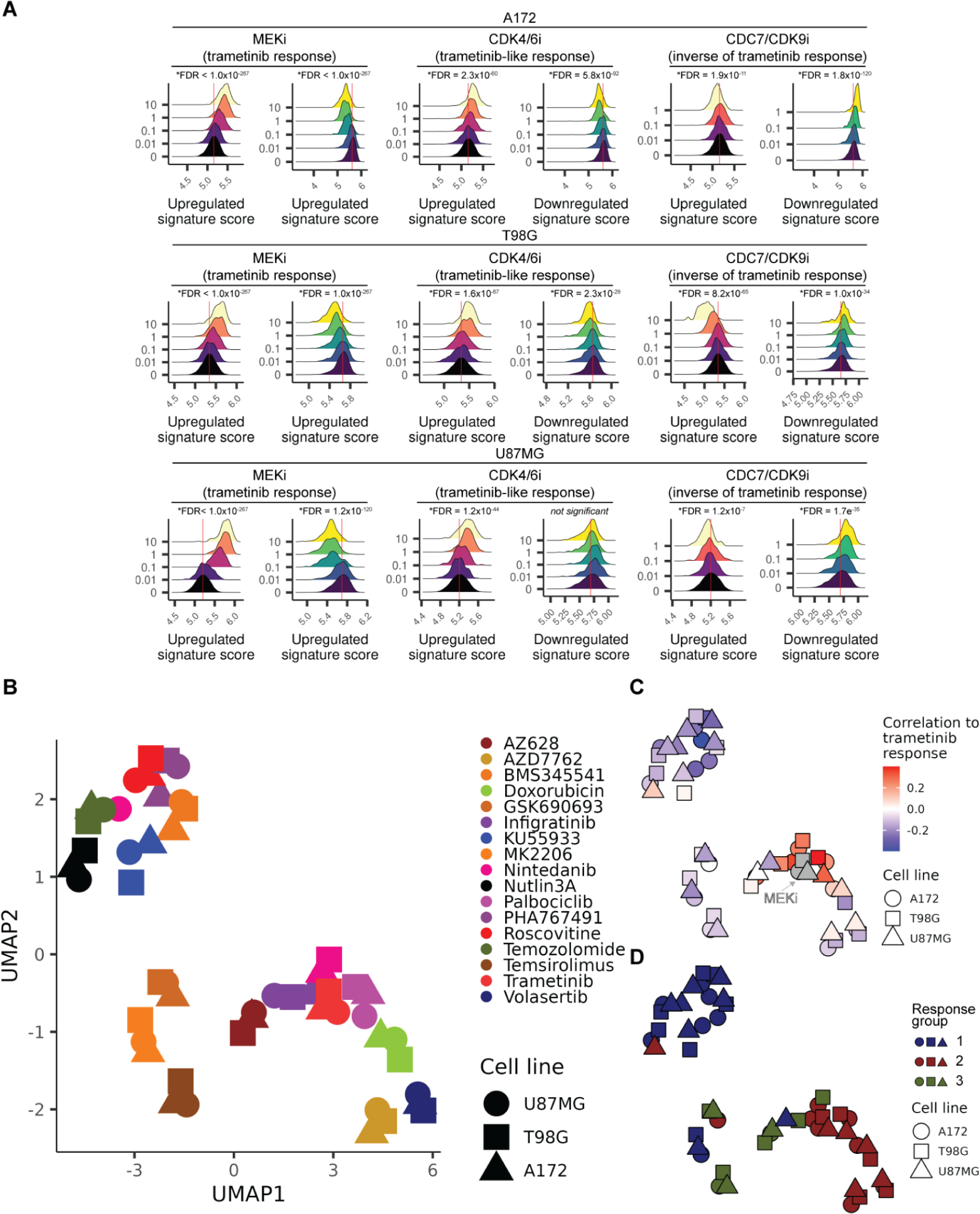
Effect and similarity of CDK4/6, CDC7/CDK9 and additional kinase inhibition on the trametinib induced compensatory program. **A**) Density plots of upregulated and downregulated signature scores of cells treated with the MEK inhibitor trametinib (MEKi), the CDK4/6 inhibitor palbociclib (CDK4/6i) or the CDC7 inhibitor PHA767491 (CDC7i) for the three glioblastoma cell lines. The 10 µM dose for PHA767491 exposed A172 and U87MG cells have been removed due to low recovery of cells for those exposures. Red vertical lines denote the mean signature expression of vehicle exposed cells. *FDR < 0.05. **B**) UMAP embeddings summarizing the pair-wise correlation of all specified single exposures across genes that compose the compensatory program enacted by RTK pathway inhibition. Shapes refer to individual cell lines. **C**) UMAP as in **B,** colored by the correlation of each single exposure to cells exposed to trametinib alone across genes that compose the compensatory program enacted by RTK pathway inhibition. **D**) UMAP as in **B** colored by response group clusters identified by Leiden-based community detection.

**Supplementary Figure 9.**
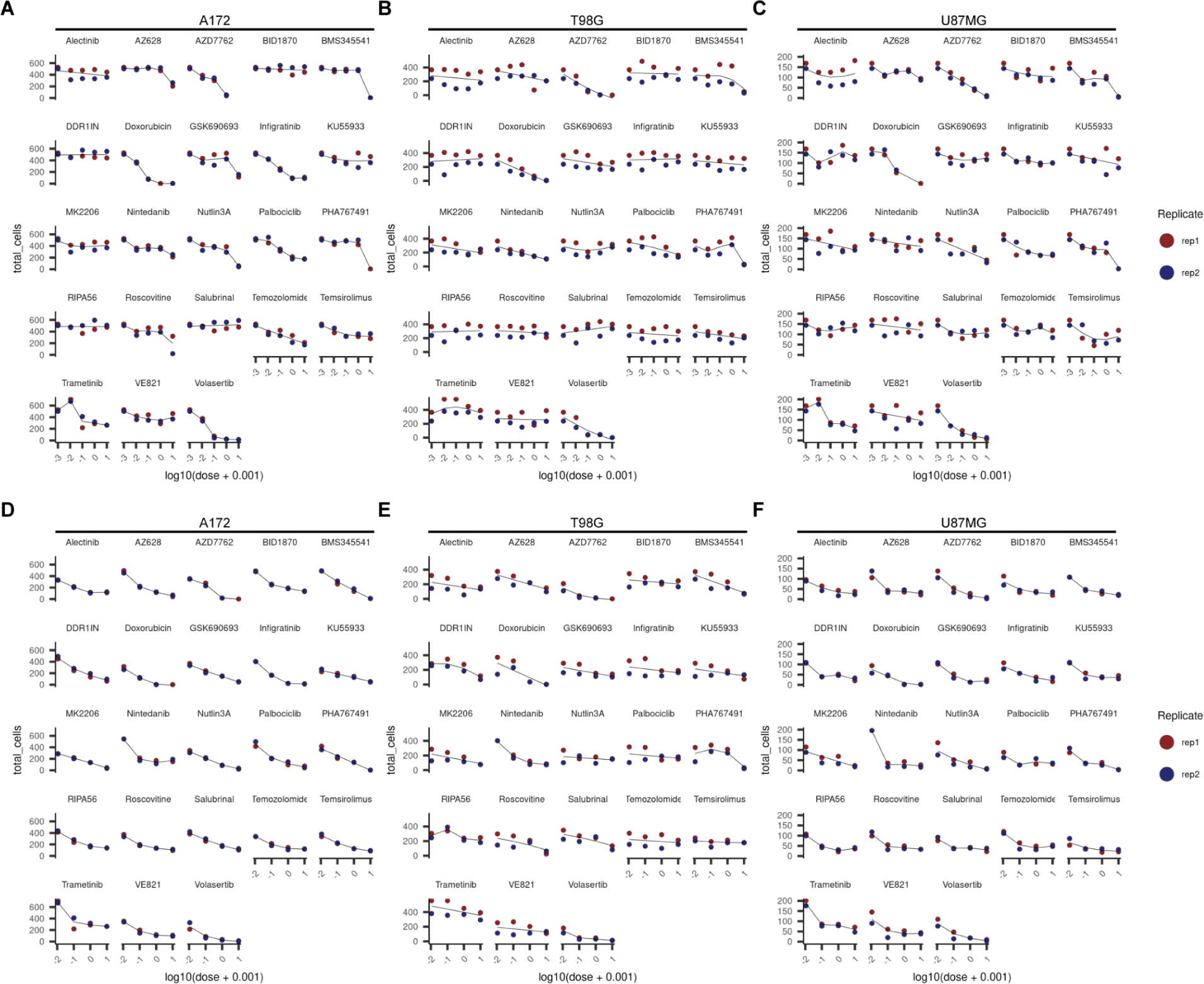
Summaries of the effect of single and combinatorial chemical exposure on the expression of conserved S1 and S2 signature genes and cellular viability. **A-C**) Cell count viability estimates derived from sci-Plex hash labels for each treatment and dose across A172 (**B**), T98G (**C**), and U87MG (**D**) GBM cells. **D-F**) Cell count viability estimates as in A-C for combinatorial trametinibn exposure across A172 (**D**), T98G (**E**), and U87MG (**F**) GBM cells.

## Supplementary Tables

**Supplementary table 1:**
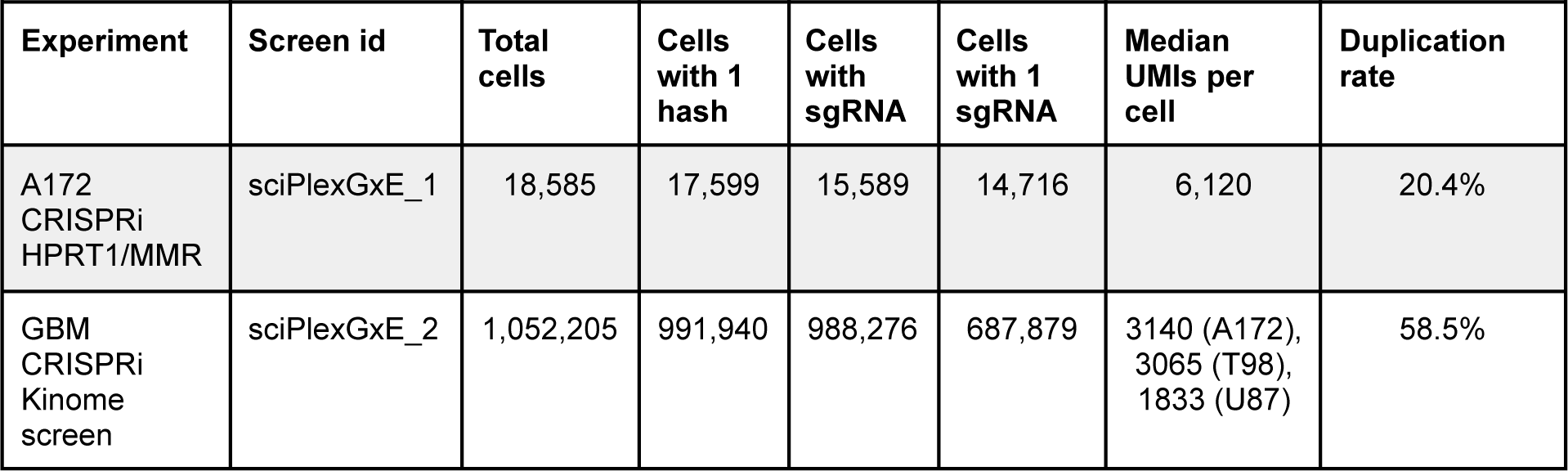
Summary of sci-Plex-GxE experiments in this study. Screen id refers to the experiment identifier in the NCBI GEO submission of our dataset.

**Supplementary table 2:**
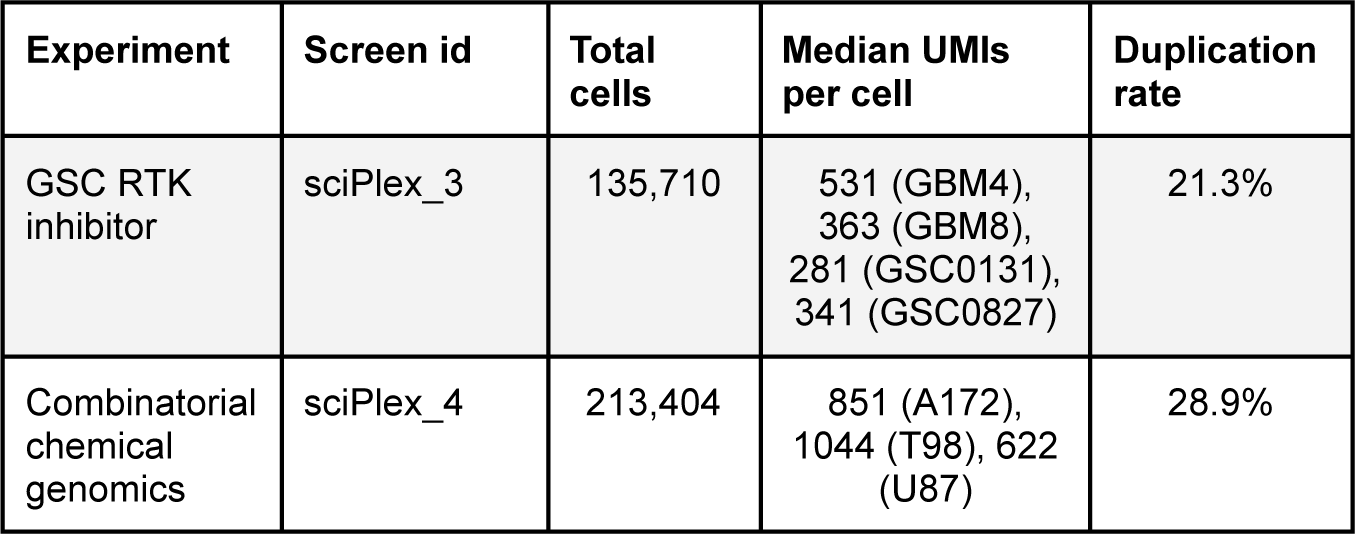
Summary of sci-Plex chemical transcriptomics experiments in this study. Screen id refers to the experiment identifier in the NCBI GEO submission of our dataset.

**Supplementary table 3:**
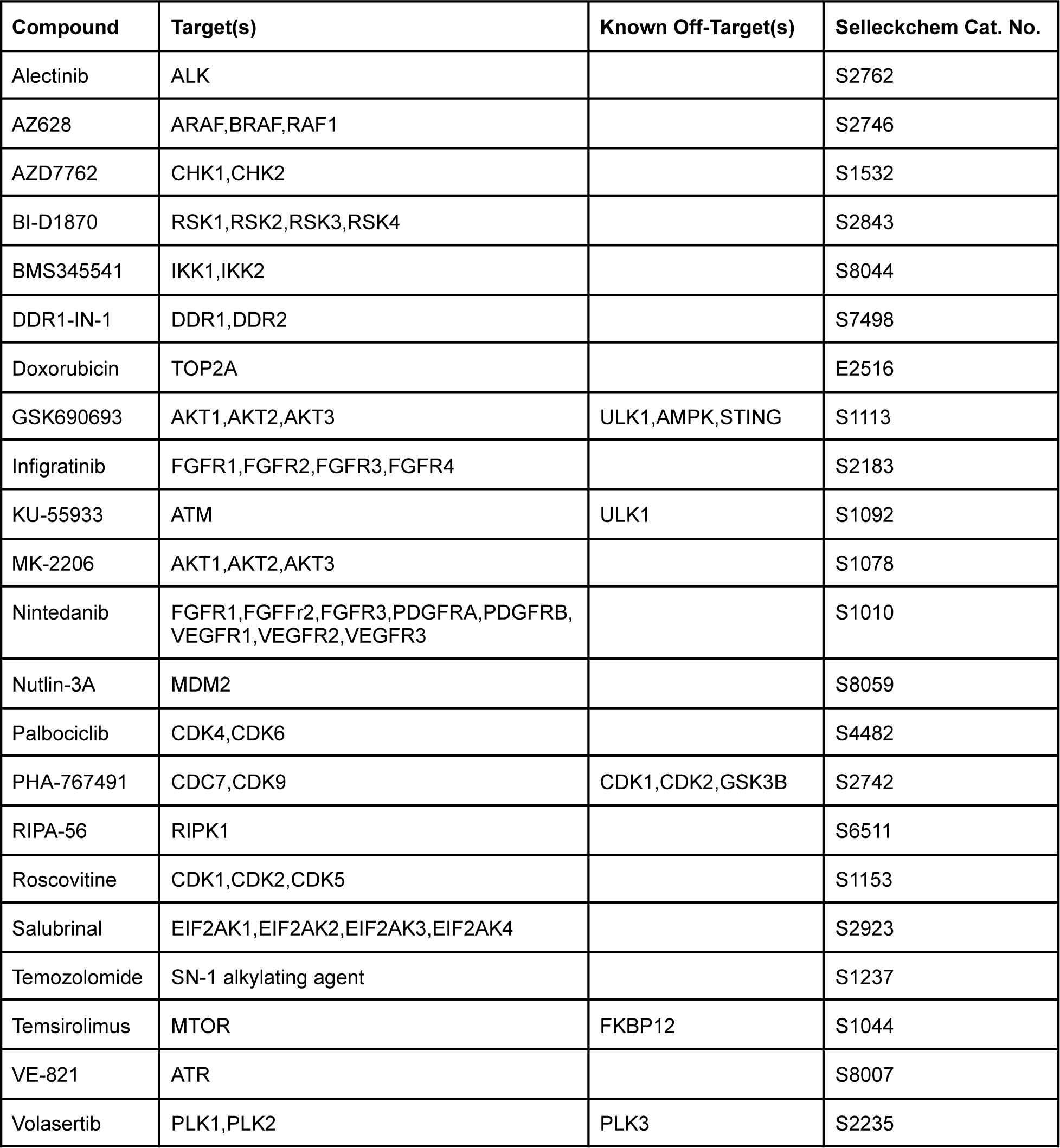
Compounds used in our validation chemical genomics screen.

## Additional Supplementary Files

Supplementary File 1: Contains the results of differential gene expression testing as a function of genotypes across TMZ exposed A172 cells.

Supplementary File 2: Contains the results of gene module analysis of the results from differential gene expression testing as a function of genotypes across TMZ exposed A172 cells (Supplementary File 1), the results from differential gene expression testing as a function of TMZ dose in NTC cells and inferred TC50 calculations.

Supplementary File 3: Contains the results of differential gene expression testing as a function of RTK pathway inhibitor dose in unperturbed cells.

Supplementary File 4: Contains the top kinase perturbations that modulate the adaptive program across the 3 GBM lines used in this study.

Supplementary File 5: Contains the sgRNA protospacer sequences used in this study.

## Notes

### Summary of Updates

Figure 2 revised, captions revised.

## References

1. Stockwell, B.R. (2000). Chemical genetics: ligand-based discovery of gene function. Nat. Rev. Genet. 1, 116–125.

2. Wood, A.J., Lo, T.-W., Zeitler, B., Pickle, C.S., Ralston, E.J., Lee, A.H., Amora, R., Miller, J.C., Leung, E., Meng, X., et al. (2011). Targeted genome editing across species using ZFNs and TALENs. Science 333, 307.

3. Wang, T., Birsoy, K., Hughes, N.W., Krupczak, K.M., Post, Y., Wei, J.J., Lander, E.S., and Sabatini, D.M. (2015). Identification and characterization of essential genes in the human genome. Science 350, 1096–1101.

4. Hart, T., Chandrashekhar, M., Aregger, M., Steinhart, Z., Brown, K.R., MacLeod, G., Mis, M., Zimmermann, M., Fradet-Turcotte, A., Sun, S., et al. (2015). High-Resolution CRISPR Screens Reveal Fitness Genes and Genotype-Specific Cancer Liabilities. Cell 163, 1515–1526.

5. Adamson, B., Norman, T.M., Jost, M., Cho, M.Y., Nuñez, J.K., Chen, Y., Villalta, J.E., Gilbert, L.A., Horlbeck, M.A., Hein, M.Y., et al. (2016). A Multiplexed Single-Cell CRISPR Screening Platform Enables Systematic Dissection of the Unfolded Protein Response. Cell 167, 1867–1882.e21.

6. Dixit, A., Parnas, O., Li, B., Chen, J., Fulco, C.P., Jerby-Arnon, L., Marjanovic, N.D., Dionne, D., Burks, T., Raychowdhury, R., et al. (2016). Perturb-Seq: Dissecting Molecular Circuits with Scalable Single-Cell RNA Profiling of Pooled Genetic Screens. Cell 167, 1853–1866.e17.

7. Jaitin, D.A., Weiner, A., Yofe, I., Lara-Astiaso, D., Keren-Shaul, H., David, E., Salame, T.M., Tanay, A., van Oudenaarden, A., and Amit, I. (2016). Dissecting Immune Circuits by Linking CRISPR-Pooled Screens with Single-Cell RNA-Seq. Cell 167, 1883–1896.e15.

8. Datlinger, P., Rendeiro, A.F., Schmidl, C., Krausgruber, T., Traxler, P., Klughammer, J., Schuster, L.C., Kuchler, A., Alpar, D., and Bock, C. (2017). Pooled CRISPR screening with single-cell transcriptome readout. Nat. Methods 14, 297–301.

9. Frangieh, C.J., Melms, J.C., Thakore, P.I., Geiger-Schuller, K.R., Ho, P., Luoma, A.M., Cleary, B., Jerby-Arnon, L., Malu, S., Cuoco, M.S., et al. (2021). Multimodal pooled Perturb-CITE-seq screens in patient models define mechanisms of cancer immune evasion. Nat. Genet. 53, 332–341.

10. Norman, T.M., Horlbeck, M.A., Replogle, J.M., Ge, A.Y., Xu, A., Jost, M., Gilbert, L.A., and Weissman, J.S. (2019). Exploring genetic interaction manifolds constructed from rich single-cell phenotypes. Science 365, 786–793.

11. Replogle, J.M., Saunders, R.A., Pogson, A.N., Hussmann, J.A., Lenail, A., Guna, A., Mascibroda, L., Wagner, E.J., Adelman, K., Bonnar, J.L., et al. (2021). Mapping information-rich genotype-phenotype landscapes with genome-scale Perturb-seq. bioRxiv. 10.1101/2021.12.16.473013.

12. Cao, J., Spielmann, M., Qiu, X., Huang, X., Ibrahim, D.M., Hill, A.J., Zhang, F., Mundlos, S., Christiansen, L., Steemers, F.J., et al. (2019). The single-cell transcriptional landscape of mammalian organogenesis. Nature 566, 496–502.

13. Srivatsan, S.R., McFaline-Figueroa, J.L., Ramani, V., Saunders, L., Cao, J., Packer, J., Pliner, H.A., Jackson, D.L., Daza, R.M., Christiansen, L., et al. (2020). Massively multiplex chemical transcriptomics at single-cell resolution. Science 367, 45–51.

14. Stupp, R., Mason, W.P., van den Bent, M.J., Weller, M., Fisher, B., Taphoorn, M.J.B., Belanger, K., Brandes, A.A., Marosi, C., Bogdahn, U., et al. (2005). Radiotherapy plus Concomitant and Adjuvant Temozolomide for Glioblastoma. New England Journal of Medicine 352, 987–996. 10.1056/nejmoa043330.

15. Aquilina, G., Crescenzi, M., and Bignami, M. (1999). Mismatch repair, G 2 /M cell cycle arrest and lethality after DNA damage. Carcinogenesis 20, 2317–2326. 10.1093/carcin/20.12.2317.

16. Cejka, P. (2003). Methylation-induced G2/M arrest requires a full complement of the mismatch repair protein hMLH1. The EMBO Journal 22, 2245–2254. 10.1093/emboj/cdg216.

17. Manning, G., Whyte, D.B., Martinez, R., Hunter, T., and Sudarsanam, S. (2002). The protein kinase complement of the human genome. Science 298, 1912–1934.

18. Brennan, C.W., Verhaak, R.G.W., McKenna, A., Campos, B., Noushmehr, H., Salama, S.R., Zheng, S., Chakravarty, D., Sanborn, J.Z., Berman, S.H., et al. (2013). The somatic genomic landscape of glioblastoma. Cell 155, 462–477.

19. Körber, V., Yang, J., Barah, P., Wu, Y., Stichel, D., Gu, Z., Fletcher, M.N.C., Jones, D., Hentschel, B., Lamszus, K., et al. (2019). Evolutionary Trajectories of IDH Glioblastomas Reveal a Common Path of Early Tumorigenesis Instigated Years ahead of Initial Diagnosis. Cancer Cell 35, 692–704.e12.

20. Vivanco, I., Robins, H.I., Rohle, D., Campos, C., Grommes, C., Nghiemphu, P.L., Kubek, S., Oldrini, B., Chheda, M.G., Yannuzzi, N., et al. (2012). Differential sensitivity of glioma-versus lung cancer-specific EGFR mutations to EGFR kinase inhibitors. Cancer Discov. 2, 458–471.

21. Zawistowski, J.S., Bevill, S.M., Goulet, D.R., Stuhlmiller, T.J., Beltran, A.S., Olivares-Quintero, J.F., Singh, D., Sciaky, N., Parker, J.S., Rashid, N.U., et al. (2017). Enhancer Remodeling during Adaptive Bypass to MEK Inhibition Is Attenuated by Pharmacologic Targeting of the P-TEFb Complex. Cancer Discov. 7, 302–321.

22. Stuhlmiller, T.J., Miller, S.M., Zawistowski, J.S., Nakamura, K., Beltran, A.S., Duncan, J.S., Angus, S.P., Collins, K.A.L., Granger, D.A., Reuther, R.A., et al. (2015). Inhibition of Lapatinib-Induced Kinome Reprogramming in ERBB2-Positive Breast Cancer by Targeting BET Family Bromodomains. Cell Rep. 11, 390–404.

23. Hill, A.J., McFaline-Figueroa, J.L., Starita, L.M., Gasperini, M.J., Matreyek, K.A., Packer, J., Jackson, D., Shendure, J., and Trapnell, C. (2018). On the design of CRISPR-based single-cell molecular screens. Nature Methods 15, 271–274. 10.1038/nmeth.4604.

24. Gilbert, L.A., Larson, M.H., Morsut, L., Liu, Z., Brar, G.A., Torres, S.E., Stern-Ginossar, N., Brandman, O., Whitehead, E.H., Doudna, J.A., et al. (2013). CRISPR-mediated modular RNA-guided regulation of transcription in eukaryotes. Cell 154, 442–451.

25. Horlbeck, M.A., Gilbert, L.A., Villalta, J.E., Adamson, B., Pak, R.A., Chen, Y., Fields, A.P., Park, C.Y., Corn, J.E., Kampmann, M., et al. (2016). Compact and highly active next-generation libraries for CRISPR-mediated gene repression and activation. Elife 5. 10.7554/eLife.19760.

26. Glaab, W. (1998). Resistance to 6-thioguanine in mismatch repair-deficient human cancer cell lines correlates with an increase in induced mutations at the HPRT locus. Carcinogenesis 19, 1931–1937. 10.1093/carcin/19.11.1931.

27. Fu, D., Calvo, J.A., and Samson, L.D. (2012). Balancing repair and tolerance of DNA damage caused by alkylating agents. Nat. Rev. Cancer 12, 104–120.

28. Gaspar, N., Marshall, L., Perryman, L., Bax, D.A., Little, S.E., Viana-Pereira, M., Sharp, S.Y., Vassal, G., Pearson, A.D.J., Reis, R.M., et al. (2010). MGMT-Independent Temozolomide Resistance in Pediatric Glioblastoma Cells Associated with a PI3-Kinase–Mediated HOX/Stem Cell Gene Signature. Cancer Research 70, 9243–9252. 10.1158/0008-5472.can-10-1250.

29. Yoshioka, K.-I., Yoshioka, Y., and Hsieh, P. (2006). ATR kinase activation mediated by MutSalpha and MutLalpha in response to cytotoxic O6-methylguanine adducts. Mol. Cell 22, 501–510.

30. McFaline-Figueroa, J.L., Braun, C.J., Stanciu, M., Nagel, Z.D., Mazzucato, P., Sangaraju, D., Cerniauskas, E., Barford, K., Vargas, A., Chen, Y., et al. (2015). Minor Changes in Expression of the Mismatch Repair Protein MSH2 Exert a Major Impact on Glioblastoma Response to Temozolomide. Cancer Research 75, 3127–3138. 10.1158/0008-5472.can-14-3616.

31. Network, T.C.G.A.R., and The Cancer Genome Atlas Research Network (2008). Comprehensive genomic characterization defines human glioblastoma genes and core pathways. Nature 455, 1061–1068. 10.1038/nature07385.

32. Pazarentzos, E., and Bivona, T.G. (2015). Adaptive stress signaling in targeted cancer therapy resistance. Oncogene 34, 5599–5606.

33. Akhavan, D., Pourzia, A.L., Nourian, A.A., Williams, K.J., Nathanson, D., Babic, I., Villa, G.R., Tanaka, K., Nael, A., Yang, H., et al. (2013). De-repression of PDGFRβ transcription promotes acquired resistance to EGFR tyrosine kinase inhibitors in glioblastoma patients. Cancer Discov. 3, 534–547.

34. Duncan, J.S., Whittle, M.C., Nakamura, K., Abell, A.N., Midland, A.A., Zawistowski, J.S., Johnson, N.L., Granger, D.A., Jordan, N.V., Darr, D.B., et al. (2012). Dynamic reprogramming of the kinome in response to targeted MEK inhibition in triple-negative breast cancer. Cell 149, 307–321.

35. Wei, W., Shin, Y.S., Xue, M., Matsutani, T., Masui, K., Yang, H., Ikegami, S., Gu, Y., Herrmann, K., Johnson, D., et al. (2016). Single-Cell Phosphoproteomics Resolves Adaptive Signaling Dynamics and Informs Targeted Combination Therapy in Glioblastoma. Cancer Cell 29, 563–573.

36. Tanaka, K., Sasayama, T., Irino, Y., Takata, K., Nagashima, H., Satoh, N., Kyotani, K., Mizowaki, T., Imahori, T., Ejima, Y., et al. (2015). Compensatory glutamine metabolism promotes glioblastoma resistance to mTOR inhibitor treatment. J. Clin. Invest. 125, 1591–1602.

37. Sztal, T.E., and Stainier, D.Y.R. (2020). Transcriptional adaptation: a mechanism underlying genetic robustness. Development 147. 10.1242/dev.186452.

38. Kontarakis, Z., and Stainier, D.Y.R. (2020). Genetics in Light of Transcriptional Adaptation. Trends Genet. 36, 926–935.

39. Pérez de Castro, I., de Cárcer, G., and Malumbres, M. (2007). A census of mitotic cancer genes: new insights into tumor cell biology and cancer therapy. Carcinogenesis 28, 899–912.

40. Asquith, C.R.M., East, M.P., and Zuercher, W.J. (2019). RIOK2: straddling the kinase/ATPase line. Nat. Rev. Drug Discov. 18, 574.

41. Vanrobays, E., Gelugne, J.-P., Gleizes, P.-E., and Caizergues-Ferrer, M. (2003). Late cytoplasmic maturation of the small ribosomal subunit requires RIO proteins in Saccharomyces cerevisiae. Mol. Cell. Biol. 23, 2083–2095.

42. Lovejoy, C.A., and Cortez, D. (2009). Common mechanisms of PIKK regulation. DNA Repair 8, 1004–1008.

43. Maloveryan, A., Finta, C., Osterlund, T., and Kogerman, P. (2007). A possible role of mouse Fused (STK36) in Hedgehog signaling and Gli transcription factor regulation. J. Cell Commun. Signal. 1, 165–173.

44. Su, G.H., Bansal, R., Murphy, K.M., Montgomery, E., Yeo, C.J., Hruban, R.H., and Kern, S.E. (2001). ACVR1B (ALK4, activin receptor type 1B) gene mutations in pancreatic carcinoma. Proc. Natl. Acad. Sci. U. S. A. 98, 3254–3257.

45. Hempen, P.M., Zhang, L., Bansal, R.K., Iacobuzio-Donahue, C.A., Murphy, K.M., Maitra, A., Vogelstein, B., Whitehead, R.H., Markowitz, S.D., Willson, J.K.V., et al. (2003). Evidence of selection for clones having genetic inactivation of the activin A type II receptor (ACVR2) gene in gastrointestinal cancers. Cancer Res. 63, 994–999.

46. Hemminki, A., Tomlinson, I., Markie, D., Järvinen, H., Sistonen, P., Björkqvist, A.M., Knuutila, S., Salovaara, R., Bodmer, W., Shibata, D., et al. (1997). Localization of a susceptibility locus for Peutz-Jeghers syndrome to 19p using comparative genomic hybridization and targeted linkage analysis. Nat. Genet. 15, 87–90.

47. Ahn, Y.-H., Yang, Y., Gibbons, D.L., Creighton, C.J., Yang, F., Wistuba, I.I., Lin, W., Thilaganathan, N., Alvarez, C.A., Roybal, J., et al. (2011). Map2k4 functions as a tumor suppressor in lung adenocarcinoma and inhibits tumor cell invasion by decreasing peroxisome proliferator-activated receptor γ2 expression. Mol. Cell. Biol. 31, 4270–4285.

48. Ning, J.-F., Stanciu, M., Humphrey, M.R., Gorham, J., Wakimoto, H., Nishihara, R., Lees, J., Zou, L., Martuza, R.L., Wakimoto, H., et al. (2019). Myc targeted CDK18 promotes ATR and homologous recombination to mediate PARP inhibitor resistance in glioblastoma. Nat. Commun. 10, 2910.

49. Gil del Alcazar, C.R., Hardebeck, M.C., Mukherjee, B., Tomimatsu, N., Gao, X., Yan, J., Xie, X.-J., Bachoo, R., Li, L., Habib, A.A., et al. (2014). Inhibition of DNA double-strand break repair by the dual PI3K/mTOR inhibitor NVP-BEZ235 as a strategy for radiosensitization of glioblastoma. Clin. Cancer Res. 20, 1235–1248.

50. Jiao, W., Zhu, S., Shao, J., Zhang, X., Xu, Y., Zhang, Y., Wang, R., Zhong, Y., and Kong, D. (2022). ZSTK474 Sensitizes Glioblastoma to Temozolomide by Blocking Homologous Recombination Repair. Biomed Res. Int. 2022, 8568528.

51. Labrie, M., Brugge, J.S., Mills, G.B., and Zervantonakis, I.K. (2022). Therapy resistance: opportunities created by adaptive responses to targeted therapies in cancer. Nat. Rev. Cancer 22, 323–339.

52. Lundgren, K., Walworth, N., Booher, R., Dembski, M., Kirschner, M., and Beach, D. (1991). mik1 and wee1 cooperate in the inhibitory tyrosine phosphorylation of cdc2. Cell 64, 1111–1122.

53. Wakimoto, H., Kesari, S., Farrell, C.J., Curry, W.T., Jr, Zaupa, C., Aghi, M., Kuroda, T., Stemmer-Rachamimov, A., Shah, K., Liu, T.-C., et al. (2009). Human glioblastoma-derived cancer stem cells: establishment of invasive glioma models and treatment with oncolytic herpes simplex virus vectors. Cancer Res. 69, 3472–3481.

54. Toledo, C.M., Ding, Y., Hoellerbauer, P., Davis, R.J., Basom, R., Girard, E.J., Lee, E., Corrin, P., Hart, T., Bolouri, H., et al. (2015). Genome-wide CRISPR-Cas9 Screens Reveal Loss of Redundancy between PKMYT1 and WEE1 in Glioblastoma Stem-like Cells. Cell Rep. 13, 2425–2439.

55. Mikheeva, S.A., Mikheev, A.M., Petit, A., Beyer, R., Oxford, R.G., Khorasani, L., Maxwell, J.-P., Glackin, C.A., Wakimoto, H., González-Herrero, I., et al. (2010). TWIST1 promotes invasion through mesenchymal change in human glioblastoma. Mol. Cancer 9, 194.

56. Subramanian, A., Tamayo, P., Mootha, V.K., Mukherjee, S., Ebert, B.L., Gillette, M.A., Paulovich, A., Pomeroy, S.L., Golub, T.R., Lander, E.S., et al. (2005). Gene set enrichment analysis: a knowledge-based approach for interpreting genome-wide expression profiles. Proc. Natl. Acad. Sci. U. S. A. 102, 15545–15550.

57. Liberzon, A., Subramanian, A., Pinchback, R., Thorvaldsdóttir, H., Tamayo, P., and Mesirov, J.P. (2011). Molecular signatures database (MSigDB) 3.0. Bioinformatics 27, 1739–1740.

58. Tadesse, S., Anshabo, A.T., Portman, N., Lim, E., Tilley, W., Caldon, C.E., and Wang, S. (2020). Targeting CDK2 in cancer: challenges and opportunities for therapy. Drug Discov. Today 25, 406–413.

59. Herrera-Abreu, M.T., Palafox, M., Asghar, U., Rivas, M.A., Cutts, R.J., Garcia-Murillas, I., Pearson, A., Guzman, M., Rodriguez, O., Grueso, J., et al. (2016). Early Adaptation and Acquired Resistance to CDK4/6 Inhibition in Estrogen Receptor-Positive Breast Cancer. Cancer Res. 76, 2301–2313.

60. Mueller, S.B., and Kontos, C.D. (2016). Tie1: an orphan receptor provides context for angiopoietin-2/Tie2 signaling. J. Clin. Invest. 126, 3188–3191.

61. Lee, O.-H., Xu, J., Fueyo, J., Fuller, G.N., Aldape, K.D., Alonso, M.M., Piao, Y., Liu, T.-J., Lang, F.F., Bekele, B.N., et al. (2006). Expression of the receptor tyrosine kinase Tie2 in neoplastic glial cells is associated with integrin beta1-dependent adhesion to the extracellular matrix. Mol. Cancer Res. 4, 915–926.

62. Ishibashi, M., Toyoshima, M., Zhang, X., Hasegawa-Minato, J., Shigeta, S., Usui, T., Kemp, C.J., Grandori, C., Kitatani, K., and Yaegashi, N. (2018). Tyrosine kinase receptor TIE-1 mediates platinum resistance by promoting nucleotide excision repair in ovarian cancer. Sci. Rep. 8, 13207.

63. Devaiah, B.N., Lewis, B.A., Cherman, N., Hewitt, M.C., Albrecht, B.K., Robey, P.G., Ozato, K., Sims, R.J., 3rd, and Singer, D.S. (2012). BRD4 is an atypical kinase that phosphorylates serine2 of the RNA polymerase II carboxy-terminal domain. Proc. Natl. Acad. Sci. U. S. A. 109, 6927–6932.

64. Metz, K.S., Deoudes, E.M., Berginski, M.E., Jimenez-Ruiz, I., Aksoy, B.A., Hammerbacher, J., Gomez, S.M., and Phanstiel, D.H. (2018). Coral: Clear and Customizable Visualization of Human Kinome Data. Cell Syst 7, 347–350.e1.

65. Labib, K. (2010). How do Cdc7 and cyclin-dependent kinases trigger the initiation of chromosome replication in eukaryotic cells? Genes Dev. 24, 1208–1219.

66. Price, D.H. (2000). P-TEFb, a cyclin-dependent kinase controlling elongation by RNA polymerase II. Mol. Cell. Biol. 20, 2629–2634.

67. Cai, D., Latham, V.M., Jr, Zhang, X., and Shapiro, G.I. (2006). Combined depletion of cell cycle and transcriptional cyclin-dependent kinase activities induces apoptosis in cancer cells. Cancer Res. 66, 9270–9280.

68. Storch, K., and Cordes, N. (2016). The impact of CDK9 on radiosensitivity, DNA damage repair and cell cycling of HNSCC cancer cells. Int. J. Oncol. 48, 191–198.

69. Brown, W.S., McDonald, P.C., Nemirovsky, O., Awrey, S., Chafe, S.C., Schaeffer, D.F., Li, J., Renouf, D.J., Stanger, B.Z., and Dedhar, S. (2020). Overcoming Adaptive Resistance to KRAS and MEK Inhibitors by Co-targeting mTORC1/2 Complexes in Pancreatic Cancer. Cell Rep Med 1, 100131.

70. Larkin, J., Ascierto, P.A., Dréno, B., Atkinson, V., Liszkay, G., Maio, M., Mandalà, M., Demidov, L., Stroyakovskiy, D., Thomas, L., et al. (2014). Combined vemurafenib and cobimetinib in BRAF-mutated melanoma. N. Engl. J. Med. 371, 1867–1876.

71. Long, G.V., Stroyakovskiy, D., Gogas, H., Levchenko, E., de Braud, F., Larkin, J., Garbe, C., Jouary, T., Hauschild, A., Grob, J.J., et al. (2014). Combined BRAF and MEK inhibition versus BRAF inhibition alone in melanoma. N. Engl. J. Med. 371, 1877–1888.

72. Robert, C., Karaszewska, B., Schachter, J., Rutkowski, P., Mackiewicz, A., Stroiakovski, D., Lichinitser, M., Dummer, R., Grange, F., Mortier, L., et al. (2015). Improved overall survival in melanoma with combined dabrafenib and trametinib. N. Engl. J. Med. 372, 30–39.

73. Hanahan, D., and Weinberg, R.A. (2011). Hallmarks of cancer: the next generation. Cell 144, 646–674.

74. Tsherniak, A., Vazquez, F., Montgomery, P.G., Weir, B.A., Kryukov, G., Cowley, G.S., Gill, S., Harrington, W.F., Pantel, S., Krill-Burger, J.M., et al. (2017). Defining a Cancer Dependency Map. Cell 170, 564–576.e16.

75. Subramanian, A., Narayan, R., Corsello, S.M., Peck, D.D., Natoli, T.E., Lu, X., Gould, J., Davis, J.F., Tubelli, A.A., Asiedu, J.K., et al. (2017). A Next Generation Connectivity Map: L1000 Platform and the First 1,000,000 Profiles. Cell 171, 1437–1452.e17.

76. Gottifredi, V., Karni-Schmidt, O., Shieh, S.S., and Prives, C. (2001). p53 down-regulates CHK1 through p21 and the retinoblastoma protein. Mol. Cell. Biol. 21, 1066–1076.

77. Son, M.J., Woolard, K., Nam, D.-H., Lee, J., and Fine, H.A. (2009). SSEA-1 is an enrichment marker for tumor-initiating cells in human glioblastoma. Cell Stem Cell 4, 440–452.

78. Wang, X., Prager, B.C., Wu, Q., Kim, L.J.Y., Gimple, R.C., Shi, Y., Yang, K., Morton, A.R., Zhou, W., Zhu, Z., et al. (2018). Reciprocal Signaling between Glioblastoma Stem Cells and Differentiated Tumor Cells Promotes Malignant Progression. Cell Stem Cell 22, 514–528.e5.

79. Chen, B., Gilbert, L.A., Cimini, B.A., Schnitzbauer, J., Zhang, W., Li, G.-W., Park, J., Blackburn, E.H., Weissman, J.S., Qi, L.S., et al. (2013). Dynamic imaging of genomic loci in living human cells by an optimized CRISPR/Cas system. Cell 155, 1479–1491.

80. Cusanovich, D.A., Daza, R., Adey, A., Pliner, H.A., Christiansen, L., Gunderson, K.L., Steemers, F.J., Trapnell, C., and Shendure, J. (2015). Multiplex single cell profiling of chromatin accessibility by combinatorial cellular indexing. Science 348, 910–914.

81. Dobin, A., Davis, C.A., Schlesinger, F., Drenkow, J., Zaleski, C., Jha, S., Batut, P., Chaisson, M., and Gingeras, T.R. (2013). STAR: ultrafast universal RNA-seq aligner. Bioinformatics 29, 15–21.

82. Quinlan, A.R., and Hall, I.M. (2010). BEDTools: a flexible suite of utilities for comparing genomic features. Bioinformatics 26, 841–842.

83. Wolock, S.L., Lopez, R., and Klein, A.M. (2019). Scrublet: Computational Identification of Cell Doublets in Single-Cell Transcriptomic Data. Cell Syst 8, 281–291.e9.

84. Traag, V.A., Waltman, L., and van Eck, N.J. (2019). From Louvain to Leiden: guaranteeing well-connected communities. Sci. Rep. 9, 5233.

85. Väremo, L., Nielsen, J., and Nookaew, I. (2013). Enriching the gene set analysis of genome-wide data by incorporating directionality of gene expression and combining statistical hypotheses and methods. Nucleic Acids Res. 41, 4378–4391.

86. Liberzon, A., Birger, C., Thorvaldsdóttir, H., Ghandi, M., Mesirov, J.P., and Tamayo, P. (2015). The Molecular Signatures Database (MSigDB) hallmark gene set collection. Cell Syst 1, 417–425.

87. Ritz, C., Baty, F., Streibig, J.C., and Gerhard, D. (2015). Dose-Response Analysis Using R. PLoS One 10, e0146021.

88. Tirosh, I., Izar, B., Prakadan, S.M., Wadsworth, M.H., 2nd, Treacy, D., Trombetta, J.J., Rotem, A., Rodman, C., Lian, C., Murphy, G., et al. (2016). Dissecting the multicellular ecosystem of metastatic melanoma by single-cell RNA-seq. Science 352, 189–196.

