## Supplementary Figures and Materials for "Multiplex single-cell chemical genomics reveals the kinase dependence of the response to targeted therapy"

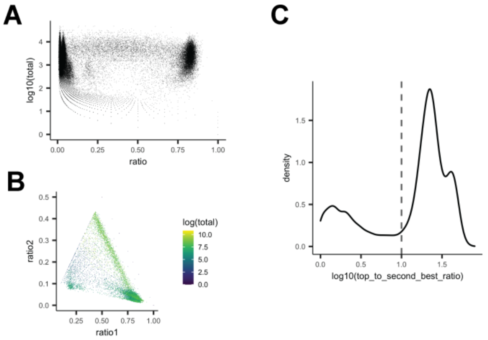


**Supplementary Figure 1. Sensitivity and specificity of sgRNA capture in the context of combinatorial indexing RNA-seq with sci-Plex-GxE.** (**A**) Plot of the log10 of the total number of sgRNA reads for a given sgRNA in a given cell as a function of the ratio of that sgRNA to all other sgRNAs in a cell (ratio). (**B**) Plot of the relationship between the proportions of the sgRNA with the highest number of reads in a cell (ratio1) vs the second most prevalent sgRNA (ratio2). (**C**) Density plot of the log10 of the top to second best ratio, the ratio between the proportion of the top most prevalent sgRNA to the second most prevalent.

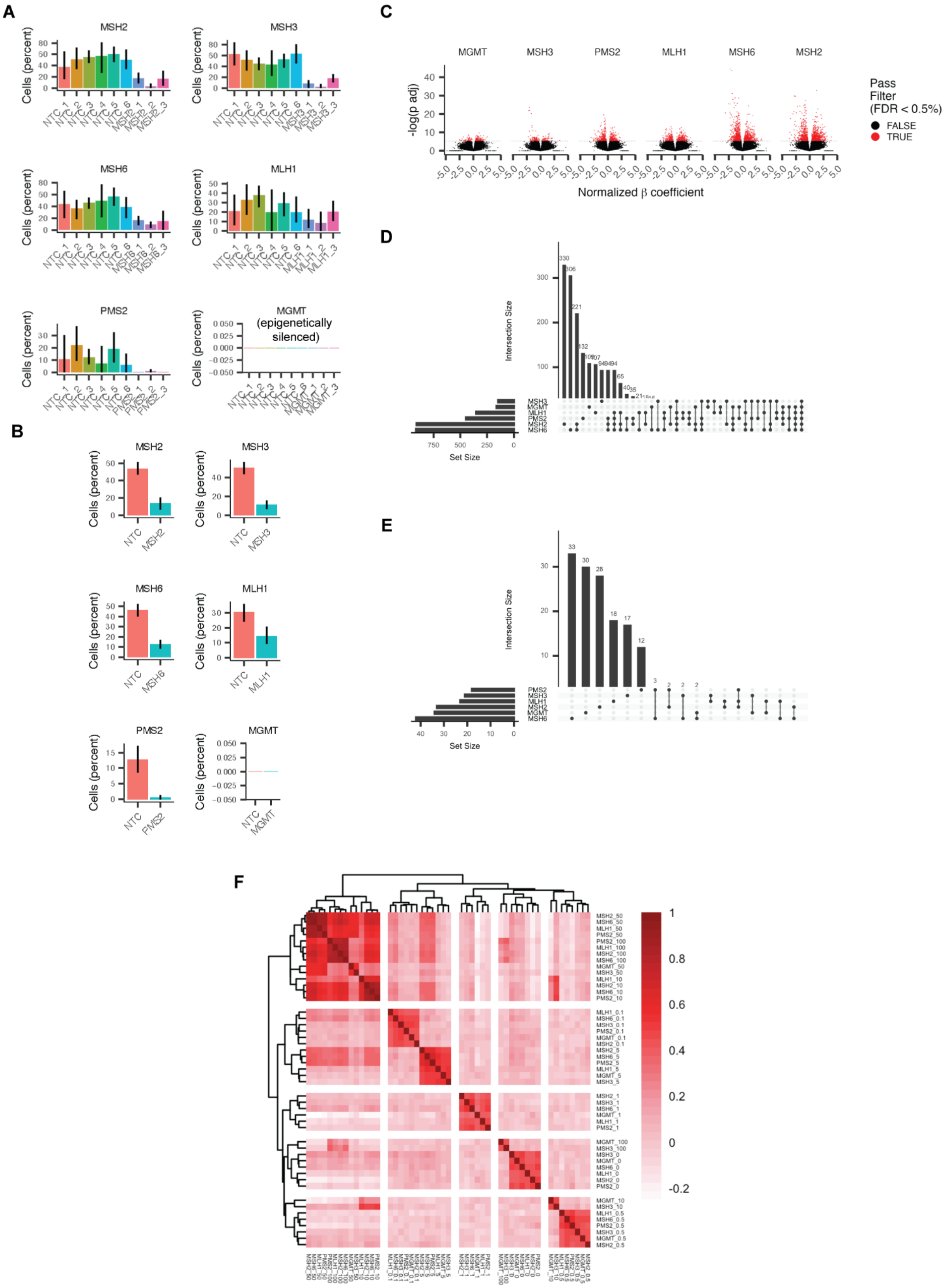

A
MSH2
MSH3
MSH6
MLH1
PMS2
MGMT (epigenetically silenced)
Cells (percent)
B
MSH2
MSH3
MSH6
MLH1
PMS2
MGMT
Cells (percent)
C
MGMT
MSH3
PMS2
MLH1
MSH6
MSH2
-log(p adj)
Normalized β coefficient
Pass Filter (FDR < 0.5%)
FALSE
TRUE
D
Intersection Size
Set Size
E
Intersection Size
Set Size
F

**Supplementary Figure 2. Effects of mismatch repair perturbation of gene expression changes induced by exposure of glioblastoma cells to the chemotherapeutic agent temozolomide.** (**A**) Bar plots of the expression level of the mismatch repair components *MSH2*, *MSH3*, *MSH6*, *MLH1*, *PMS2* and the direct repair enzyme MGMT in A172 dCas9-KRAB cells expressing individual sgRNAs against each gene or non targeting controls sgRNAs (NTC). Note that MGMT is not expressed in A172 cells due to epigenetically silencing of the *MGMT* locus by promoter methylation (citation). (**B**) Bar plots as in (**A**) across cells binned by their respective target. (**C**) Volcano plots of the relationship between statistical significance and effect size for the results of differential gene expression analysis of the effect of genetic perturbation of *MSH2*, *MSH3*, *MSH6*, *MLH1*, *PMS2,* and *MGMT* on gene expression after exposure to various doses of temozolomide. For each genotype, genetically perturbed and NTC cells were subsetted by dose, expression was log-transformed and differentially expressed genes were identified by fitting a generalized linear model of the form *expression ~ genotype*. All tests were then combined and p-values corrected for multiple hypothesis testing using Benjamini-Hochberg. Red indicates genes whose temozolomide-induced gene expression is significantly affected by genetic perturbation at FDR < 0.05. (**D-E**) Upset plots of the overlap of differentially expressed genes in the presence (**D**) or absence (**E**) of temozolomide exposure. (**F**) Correlation heatmap of the normalized effect sizes across all perturbation-dependent differentially expressed genes upon exposure to temozolomide. Rows and columns are labeled as “genotype_dose of temozolomide”. Pearson’s correlation.

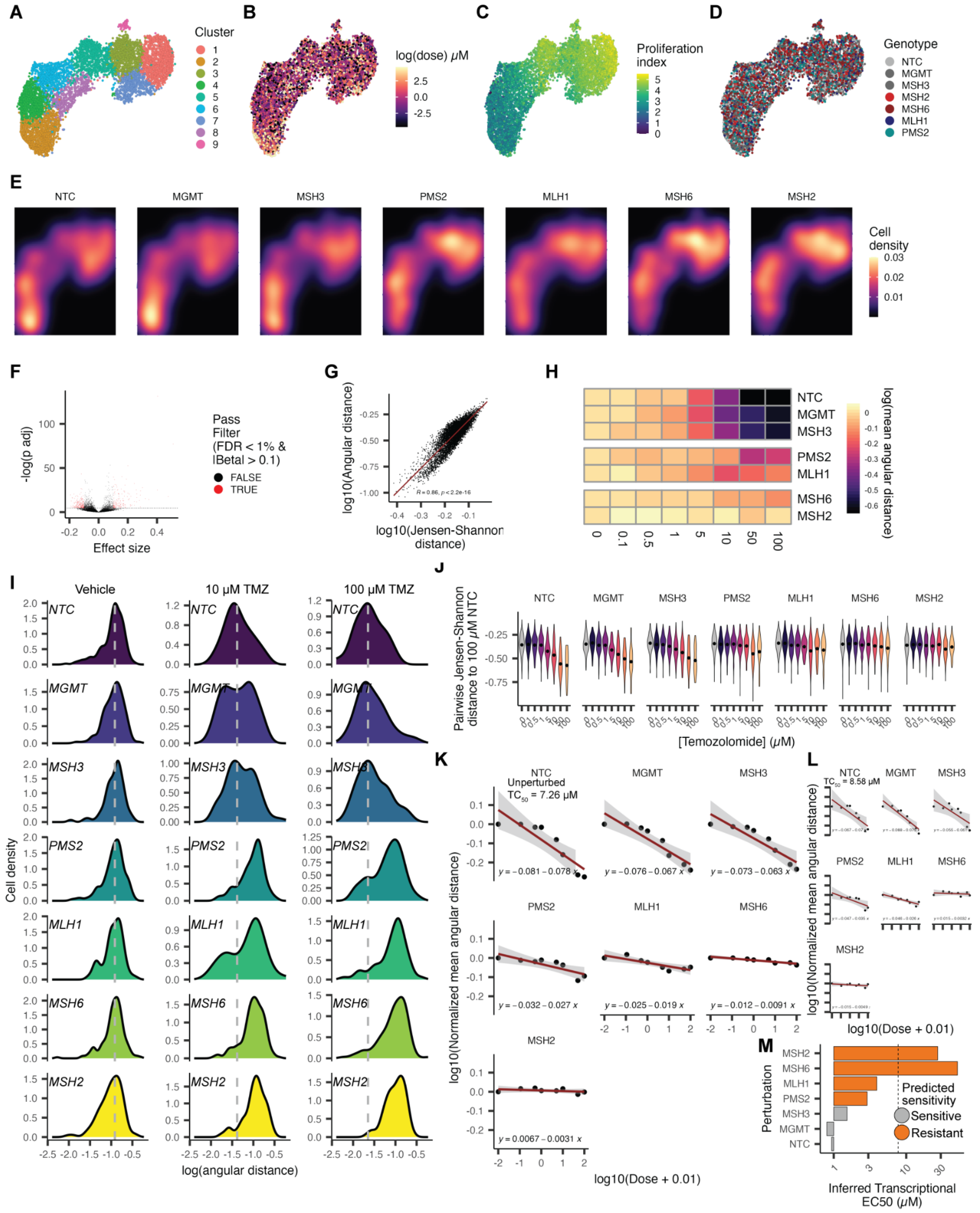

**Supplementary Figure 3. Summarizing state-level effects of temozolomide exposure as a function of genetic perturbation. A-D**) UMAP embedding of vehicle and temozolomide exposed and genetically perturbed A172 dCas9-KRAB cells. Cells are colored as a function of PCA cluster

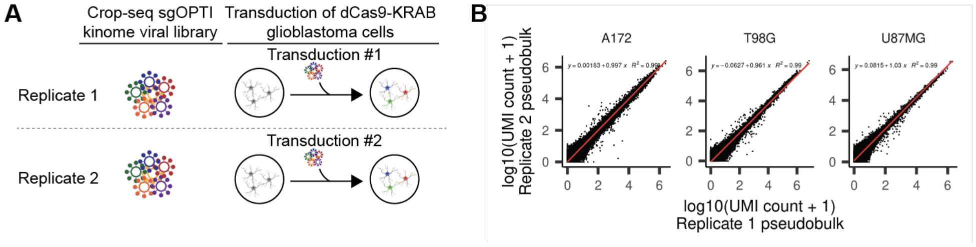


**Supplementary Figure 4. Experimental design and QC**. **A**) Schematic depicting the replicate structure of our single-cell kinome screen. **B**) Correlation between replicate screens across the three glioblastoma cell lines in our experiment.

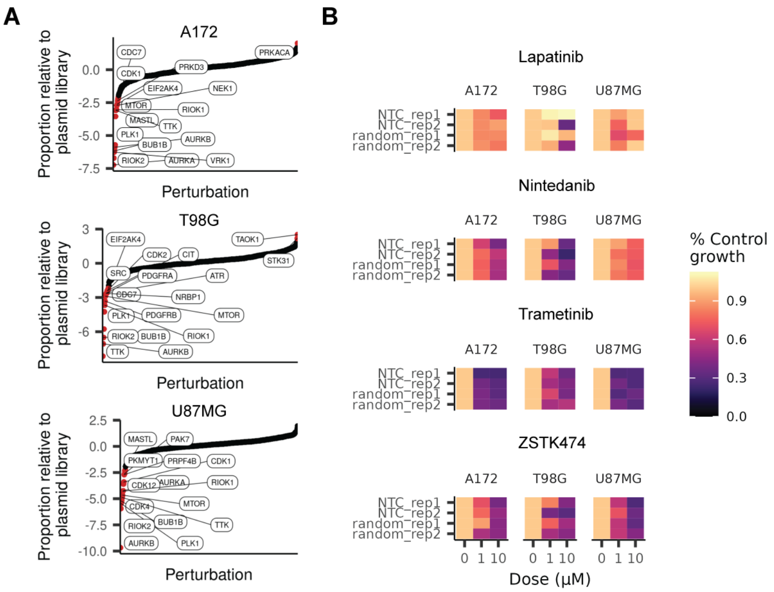


**Supplementary Figure 5: Effect of kinase perturbation and RTK pathway inhibition on GBM cell bulk viability. A.** Proportion of cells expressing sgRNAs targeting individual kinases in our screen relative to the starting proportion of sgRNAs in our CROP-seq kinome plasmid library. Labels correspond to depleted kinases over a z-score of 1. **B.** Heatmap depicting viability estimates derived from sci-Plex cell counts for unperturbed cells expressing non-targeting or random targeting control sgRNAs.

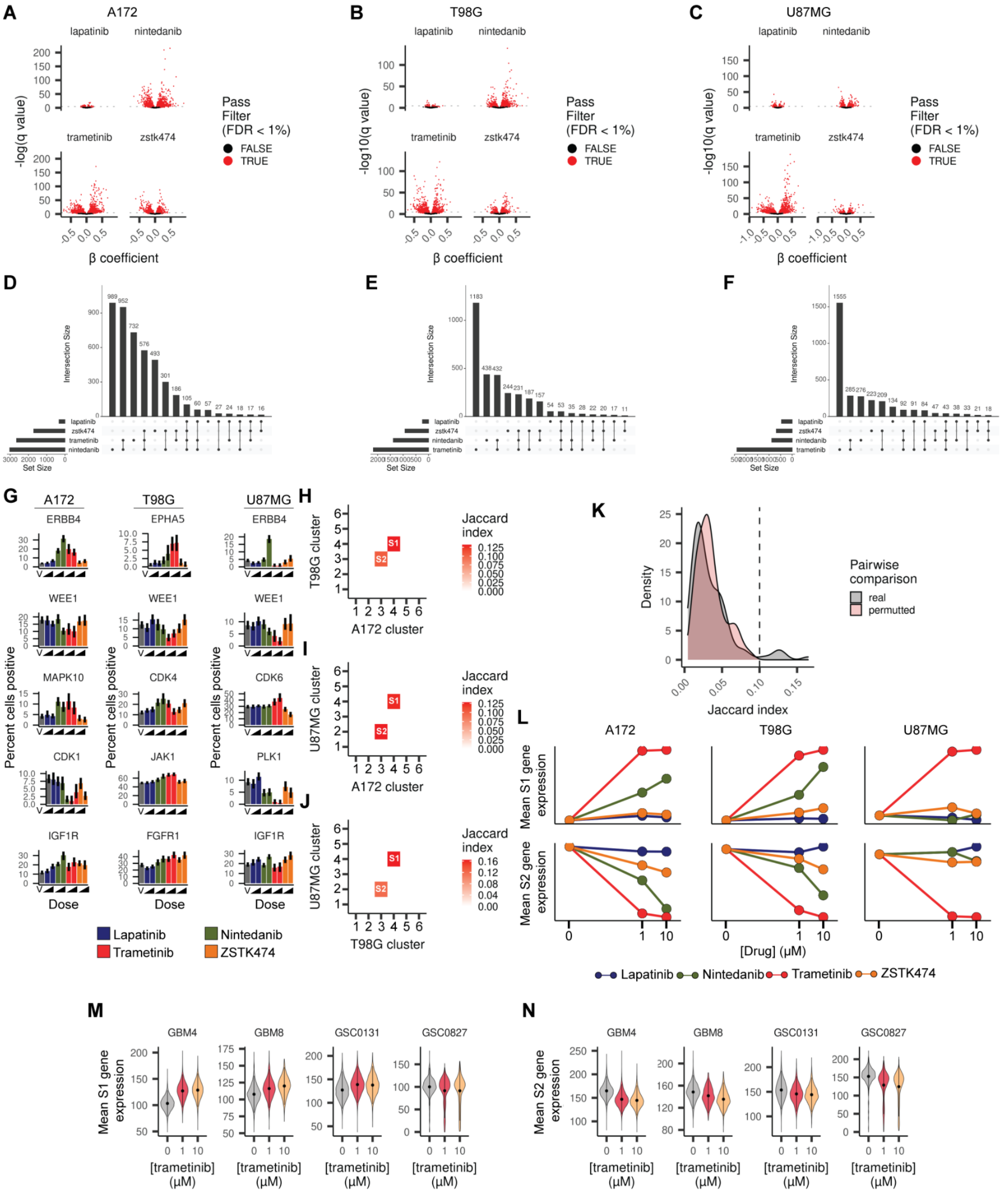

**Supplementary Figure 6. Exposure to small molecule inhibitors targeting the RTK pathway leads to dynamic rewiring of transcriptional networks in glioblastoma cells. A-C**) Volcano plots of the relationship between statistical significance and effect size of exposure to small molecules targeting the RTK pathway (lapatinib, nintedanib, trametinib, zstk474) on gene expression for A172 (**A**), T98G (**B**), and U87MG (**C**) cells. A generalized linear model of the form expression ~ log(dose) + replicate was fitted across log normalized expressions for every subset containing cells exposed to one of the 4 agents and vehicle control. Red dots correspond to genes whose expression is significantly altered by kinase perturbation at an FDR < 0.01. D-F) UpsetR plots depicting the overlap in differentially expressed genes as a function of exposure to RTK pathway targeting agents in A172 (**D**), T98G (**E**), and U87MG (**F**) GBM cells. **G**) Bar plots depicting the percent of cells positive for the specified kinase transcripts differentially expressed in at least 1 of 4 drug exposures across the 3 GBM lines in our study. **H-J**) Heatmaps depicting the top overlap of genes between cell clusters for each pair-wise combination of teh 3 GBM cell lines. Significant overlap is defined as having a Jaccard coefficient larger than 0.1 (details in **K**). **K**) Distribution of Jaccard indices for the overlap between drug-responsive gene clusters from **Fig. 4A**-**C**. Real vs. permutted refers to the Jaccard indices calculated between clusters without and with perturbation of gene ids. Based on this test, we used a Jaccard index of 0.1 or larger to collapse signature genes across cell types. **L**) Aggregate expression of conserved upregulated (S1) and downregulated (S2) gene signatures as a function of each dose, exposure, and cell type. **M-N**) Violin plots depicting the expression of conserved S1 (**M**) and S2 (**N**) signatures in trametinib exposed glioma stem cell cultures.

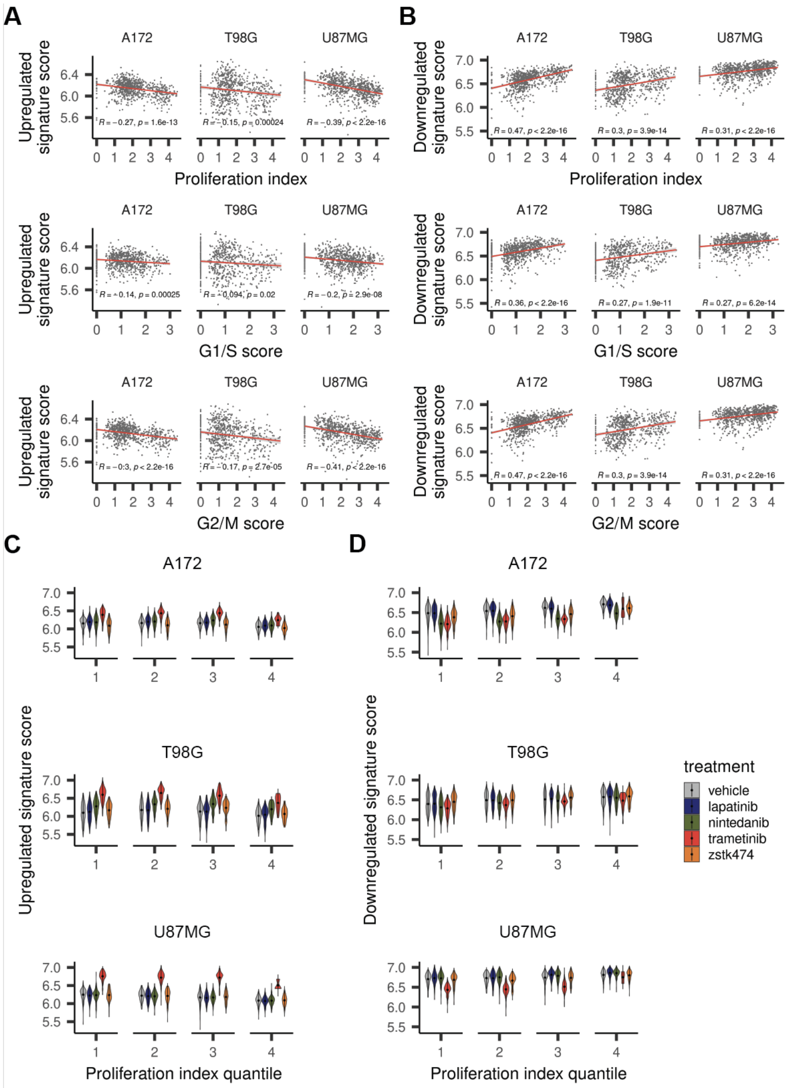


**Supplementary Figure 7. Conserved drug-dependent signatures are correlated with the cell cycle stage of a cell and by MEK kinase activity across all stages of the cell cycle. A-B**) Expression of conserved S1 (**A**) and S2 (**B**) signatures across NTC cells as a function of the aggregate expression of genes associated with proliferation (top panels), G1/S (middle panels), and G2/M (lower panels) phases of the cell cycle. **C**) Violin plots depicting the expression of S1 (**C**) and S2 (**D**) signatures as a function of treatment across cells of varying proliferation index quantiles (i.e. the x-axes of the top panel of **A** and **B** were divided into 4 equal-sized bins). Note that trametinib exposed cells have higher S1 expression and lower S2 expression across all bins compared to vehicle control.

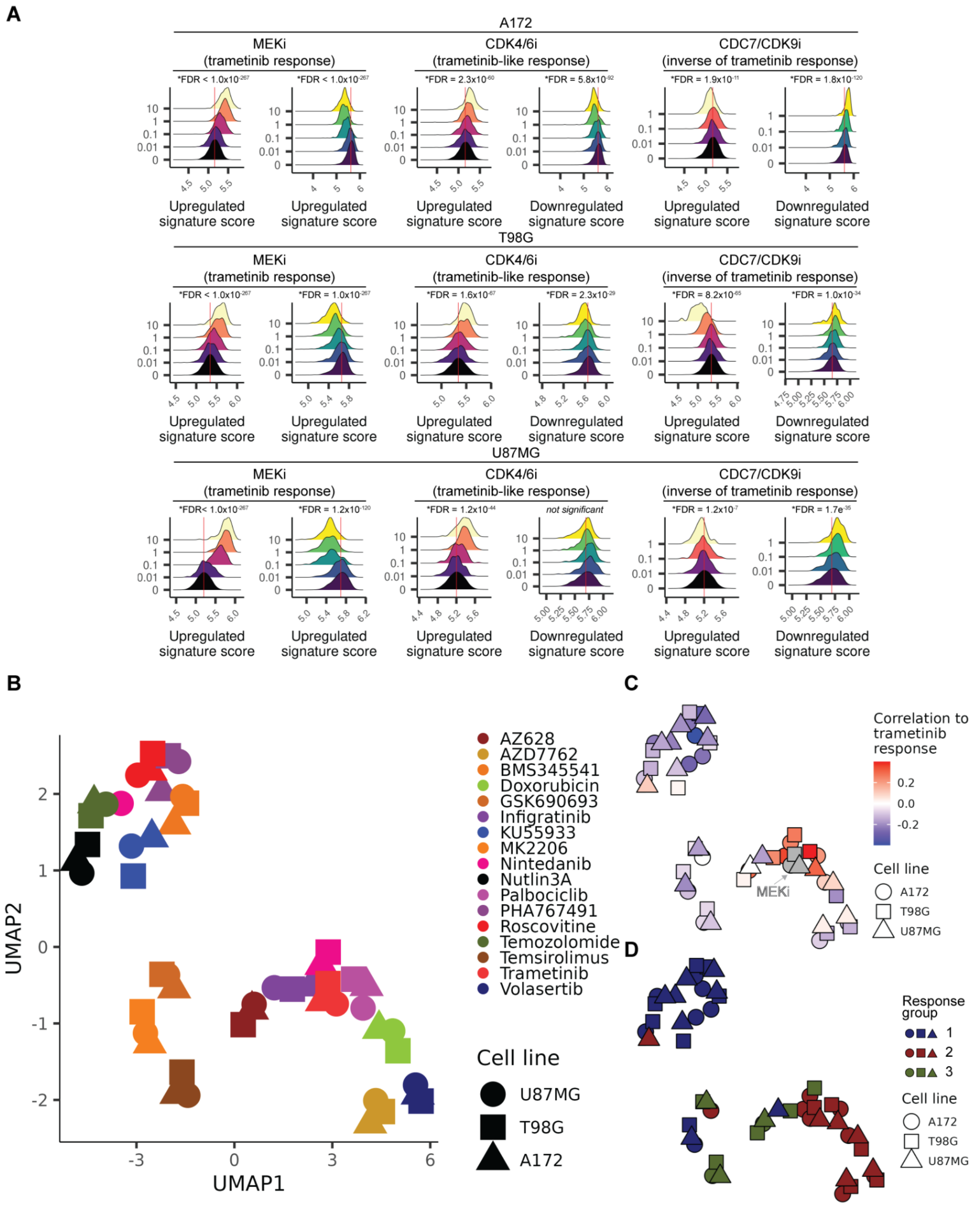


**Supplementary Figure 8. Effect and similarity of CDK4/6, CDC7/CDK9 and additional kinase inhibition on the trametinib induced compensatory program. A**) Density plots of upregulated and downregulated signature scores of cells treated with the MEK inhibitor trametinib (MEKi), the CDK4/6 inhibitor palbociclib (CDK4/6i) or the CDC7 inhibitor PHA767491 (CDC7i) for the three glioblastoma cell lines. The 10 µM dose for PHA767491 exposed A172 and U87MG cells have been removed due to low recovery of cells for those exposures. Red vertical lines denote the mean signature expression of vehicle exposed cells. *FDR < 0.05. **B**) UMAP embeddings summarizing the pair-wise correlation of all specified single exposures across genes that compose the compensatory program enacted by RTK pathway inhibition. Shapes refer to individual cell lines. **C**) UMAP as in **B,** colored by the correlation of each single exposure to cells exposed to trametinib alone across genes that compose the compensatory program enacted by RTK pathway inhibition. **D**) UMAP as in **B** colored by response group clusters identified by Leiden-based community detection.

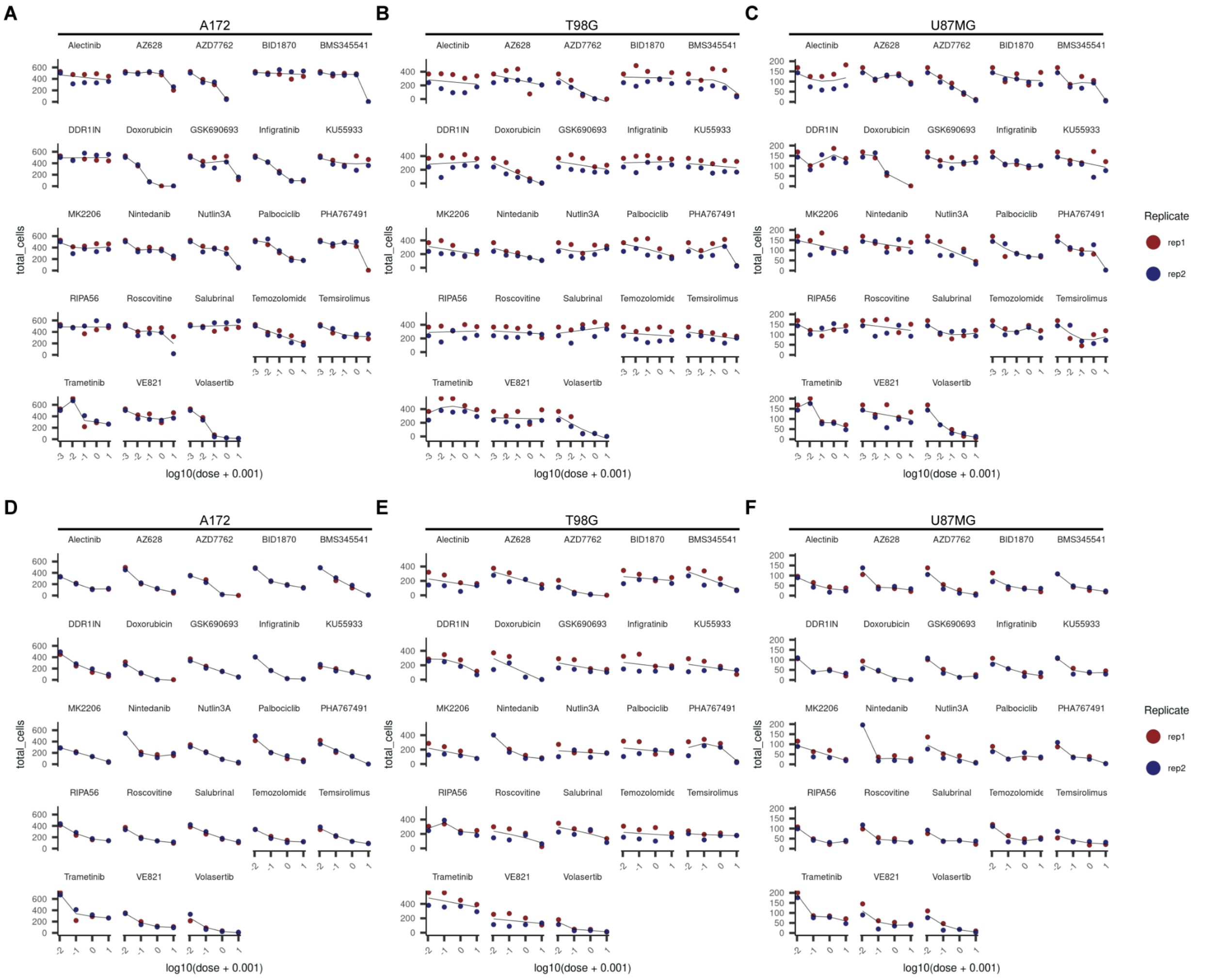


**Supplementary Figure 9. Summaries of the effect of single and combinatorial chemical exposure on the expression of conserved S1 and S2 signature genes and cellular viability. A-C**) Cell count viability estimates derived from sci-Plex hash labels for each treatment and dose across A172 (**B**), T98G (**C**), and U87MG (**D**) GBM cells. **D-F**) Cell count viability estimates as in A-C for combinatorial trametinibn exposure across A172 (**D**), T98G (**E**), and U87MG (**F**) GBM cells.

### Supplementary Tables

| Experiment | Screen id | Total cells | Cells with 1 hash | Cells with sgRNA | Cells with 1 sgRNA | Median UMIs per cell | Duplication rate |
|---|---|---|---|---|---|---|---|
| A172 CRISPRi HPRT1/MMR | sciPlexGxE_1 | 18,585 | 17,599 | 15,589 | 14,716 | 6,120 | 20.4% |
| GBM CRISPRi Kinome screen | sciPlexGxE_2 | 1,052,205 | 991,940 | 988,276 | 687,879 | 3140 (A172), 3065 (T98), 1833 (U87) | 58.5% |

**Supplementary table 1: Summary of sci-Plex-GxE experiments in this study.** Screen id refers to the experiment identifier in the NCBI GEO submission of our dataset.

| Experiment | Screen id | Total cells | Median UMIs per cell | Duplication rate |
|---|---|---|---|---|
| GSC RTK inhibitor | sciPlex_3 | 135,710 | 531 (GBM4), 363 (GBM8), 281 (GSC0131), 341 (GSC0827) | 21.3% |
| Combinatorial chemical genomics | sciPlex_4 | 213,404 | 851 (A172), 1044 (T98), 622 (U87) | 28.9% |

**Supplementary table 2: Summary of sci-Plex chemical transcriptomics experiments in this study.** Screen id refers to the experiment identifier in the NCBI GEO submission of our dataset.

| Compound | Target(s) | Known Off-Target(s) | Selleckchem Cat. No. |
|---|---|---|---|
| Alectinib | ALK | | S2762 |
| AZ628 | ARAF,BRAF,RAF1 | | S2746 |
| AZD7762 | CHK1,CHK2 | | S1532 |
| BI-D1870 | RSK1,RSK2,RSK3,RSK4 | | S2843 |
| BMS345541 | IKK1,IKK2 | | S8044 |
| DDR1-IN-1 | DDR1,DDR2 | | S7498 |
| Doxorubicin | TOP2A | | E2516 |
| GSK690693 | AKT1,AKT2,AKT3 | ULK1,AMPK,STING | S1113 |
| Infigratinib | FGFR1,FGFR2,FGFR3,FGFR4 | | S2183 |
| KU-55933 | ATM | ULK1 | S1092 |
| MK-2206 | AKT1,AKT2,AKT3 | | S1078 |
| Nintedanib | FGFR1,FGFFr2,FGFR3,PDGFRA,PDGFRB, VEGFR1,VEGFR2,VEGFR3 | | S1010 |
| Nutlin-3A | MDM2 | | S8059 |
| Palbociclib | CDK4,CDK6 | | S4482 |
| PHA-767491 | CDC7,CDK9 | CDK1,CDK2,GSK3B | S2742 |
| RIPA-56 | RIPK1 | | S6511 |
| Roscovitine | CDK1,CDK2,CDK5 | | S1153 |
| Salubrinal | EIF2AK1,EIF2AK2,EIF2AK3,EIF2AK4 | | S2923 |
| Temozolomide | SN-1 alkylating agent | | S1237 |
| Temsirolimus | MTOR | FKBP12 | S1044 |
| VE-821 | ATR | | S8007 |
| Volasertib | PLK1,PLK2 | PLK3 | S2235 |

**Supplementary table 3: Compounds used in our validation chemical genomics screen.**

**Additional Supplementary Files**

Supplementary File 1: Contains the results of differential gene expression testing as a function of genotypes across TMZ exposed A172 cells.

Supplementary File 2: Contains the results of gene module analysis of the results from differential gene expression testing as a function of genotypes across TMZ exposed A172 cells (Supplementary File 1), the results from differential gene expression testing as a function of TMZ dose in NTC cells and inferred TC50 calculations.

Supplementary File 3: Contains the results of differential gene expression testing as a function of RTK pathway inhibitor dose in unperturbed cells.

Supplementary File 4: Contains the top kinase perturbations that modulate the adaptive program across the 3 GBM lines used in this study.

Supplementary File 5: Contains the sgRNA protospacer sequences used in this study.
